# The Developmental Transcription Factor TBX3 Physically Engages with the Wnt/β-catenin Transcriptional Complex in Human Colorectal Cancer Cells to Regulate Metastasis Genes

**DOI:** 10.1101/2023.12.18.571901

**Authors:** Amaia Jauregi-Miguel, Simon Söderholm, Tamina Weiss, Anna Nordin, Valeria Ghezzi, Salome M. Brütsch, Pierfrancesco Pagella, Yorick van de Grift, Gianluca Zambanini, Jacopo Ulisse, Alessandro Mattia, Ruslan Deviatiiarov, Elena Faustini, Lavanya Moparthi, Francisca Lottersberger, Stefan Koch, Andreas E. Moor, Xiao-Feng Sun, Eleonore von Castelmur, Guojun Sheng, Claudio Cantù

## Abstract

Wnt signaling orchestrates gene expression in a plethora of processes during development and adult cell homeostasis via the action of nuclear β-catenin. Furthermore, neoplasia of the colorectal epithelium begins with aberrant Wnt/β-catenin signaling. Yet, little is known about how β-catenin generates context-specific transcriptional outcomes. We have previously identified the developmental transcription factor TBX3 as a tissue-specific component of the Wnt/β-catenin nuclear complex during mouse forelimb development. In this study, we show that TBX3 is present and functionally active in human colorectal cancers. TBX3’s genomic binding pattern suggests a regulatory role that broadly coincides with that of Wnt/β-catenin. Moreover, proteomics proximity labelling indicated that, during Wnt pathway activation, TBX3 is vicinal to several protein partners, including the transcription factors TCF/LEF and chromatin remodeling complexes which are usually found at Wnt responsive elements. Sequence and structure analysis revealed that TBX3 possesses an exposed Asp-Pro-Phe (NPF) motif predicted by AlphaFold2 Multimer to mediate direct interactions with several Wnt-activated TBX3 partners. Deletion of NPF abrogates TBX3 proximity to these partners and its ability to modulate Wnt-dependent transcription. TBX3 emerges as a key modulator of the oncogenic activity of Wnt/β-catenin in colorectal cancer, and its mechanism of action exposes a novel druggable protein-interaction surface.

## Introduction

As a highly conserved cell-to-cell communication mechanism, the Wnt/β-catenin signaling pathway coordinates a wide spectrum of processes by activating the expression of Wnt target genes ^1^. Wnt/β-catenin assumes a crucial role in nearly all facets of embryonic development and adult stem cell homeostasis, spanning pluripotency, proliferation, and differentiation ^2^. Its aberrant activation has been linked to many diseases such as developmental irregularities and various severe forms of cancer ^3^. A prime example of a disease largely driven by uncontrolled activation of the Wnt/β-catenin signaling pathway is colorectal cancer (CRC), ranking among the most lethal malignancies in humans ^4^. However, while much work has been dedicated to uncovering effective therapeutics to block oncogenic Wnt signaling, such interventions have been proven to be challenging because of the ubiquitous activity of Wnt signaling throughout the adult body and the difficulty in finding a suitable molecular target ^5–8^. Furthermore, aiming to intervene with β-catenin, the central regulator of this pathway, has presented challenges due to the rigorous control by a complex network of feedback mechanisms ^1,9,10^.

Mechanistically, the binding of extracellular WNT ligands to their receptors is transduced into an intracellular biochemical cascade that culminates in the accumulation of the β-catenin protein in the nucleus ^11^. Nuclear β-catenin binds the TCF/LEF transcription factors that are already positioned at Wnt Responsive Elements (WRE) on the chromatin together with transcriptional cofactors that include BCL9, its paralog BCL9L, PYGO1/2 ^12–14^, the ChiLS [Chip/LDB (LIM domain-binding protein) and a tetramer of SSDP (single-stranded DNA-binding protein, also known as SSBP) ^15^] along with the members of the SWI/SNF (BAF) complex and the RNA polymerase II associated factors ^16,17^.

However, a model in which a universal Wnt/β-catenin transcriptional complex drives the expression of Wnt target genes fails to explain a growing body of in vivo evidence. For instance, contrasting the notion of constitutive requirement for BCL9 and PYGO proteins for β-catenin nuclear activity, deletion of the genes encoding for these cofactors, in the mouse, causes later embryonic lethality and phenotypes that are significantly different than those caused by mutations in Ctnnb1, the gene encoding for β-catenin ^18–21^. Analogously, different cancer contexts characterized by high Wnt signaling display different requirements for these factors. For instance, both BCL9 and PYGO proteins play an important role in promoting epithelial-to-mesenchymal transition (EMT) in breast tumors ^22–25^. In contrast, BCL9 is involved in neoplastic lesions and EMT of the colorectal epithelium whereas PYGO only seems to play a minor or no contribution ^26–29^. Hence, the context-specific elements of the Wnt transcriptional apparatus are still unclear. We consider it imperative to delve into their discovery in order to precisely tailor our approach to modulating Wnt signaling in the context of colorectal cancer ^30^.

We have previously identified the developmental transcription factor TBX3 as a participant of the Wnt-mediated transcriptional regulation, serving as a PYGO-independent cofactor for the BCL9/9L adaptor proteins in developing mouse forelimbs ^26^. In this context, TBX3 displayed a bimodal behaviour: while some TBX3 genomic binding sites were independent from Wnt/β-catenin and were accompanied by the classical T-box consensus sequence on the DNA, more than two thirds of its genome-wide distribution displayed TCF/LEF binding motifs ^26^. This, and the fact that in these regions TBX3 binding to chromatin was abrogated when BCL9/9L and/or β-catenin were mutated ^26^, was evidence for TBX3 association to WREs as co-factor rather than a DNA-binding protein.

Here, we show that the role of TBX3 as Wnt signaling component goes beyond mouse embryonic development. First, TBX3 is expressed in cells derived from CRC patients, as well as in datasets obtained from several cancer cell lines. Second, Wnt-driven CRC cells possess a distinct and extensive TBX3 genome-wide binding profile, which overlaps greatly with that of β-catenin. In these cells, TBX3 controls the expression of genes that mediate metastatic transformation. Third, in CRC cells, TBX3 associates to the Wnt/β-catenin transcriptional complex via protein-protein interactions that are mediated by an evolutionarily conserved Arg-Pro-Phe (NPF) tri-aminoacidic motif at the C-terminus of the T-box binding domain. Deletion of the NPF resulted in the loss of TBX3 protein interactions and abrogated the modulation of Wnt/β-catenin-mediated transcription. This work therefore establishes TBX3 as a relevant player in CRC cell behavior and reveals a novel interaction surface that could be targeted for future CRC molecular therapy.

## Results

### TBX3 is expressed in human colorectal cancer cells together with Wnt target genes and Wnt pathway components

We have previously identified TBX3 as a tissue-specific physical component of the Wnt/β-catenin transcriptional complex during mouse forelimb development ^26^. Moreover, while we showed that TBX3 overexpression in CRC cells was capable of enhancing their metastatic behaviour, our former study did not investigate whether TBX3 was endogenously expressed in human CRC cells. Gene expression analysis of patient-derived human cancer cell lines obtained from the FANTOM 5 Consortium ^31^ showed that *TBX3* is indeed expressed in several tumor types of endo- and mesodermal origin, including tumors from the small intestine, the colon, the rectum, and the breast (Figure 1A). It is likely that *TBX3* transcript levels are elevated in these types of tumors as a consequence of high Wnt/β-catenin signaling, as TBX3 is a bona fide Wnt target gene ^32^. However, *TBX3* expression only partially overlapped with that of other Wnt target genes such as *AXIN2* and *LGR5* (Figure 1A), suggesting varying levels of Wnt activation across tumor types, and mechanisms of uncoupled regulation between *TBX3* and other targets depending on the context ^33^. When considering the family of TBX transcription factors, TBX3 exhibited the highest expression level across the tumor cell types considered (Figure 1B), supporting the dual nature of this protein as not only regulator of developmental processes, but also of tumorigenesis ^34^.

**Figure 1.**
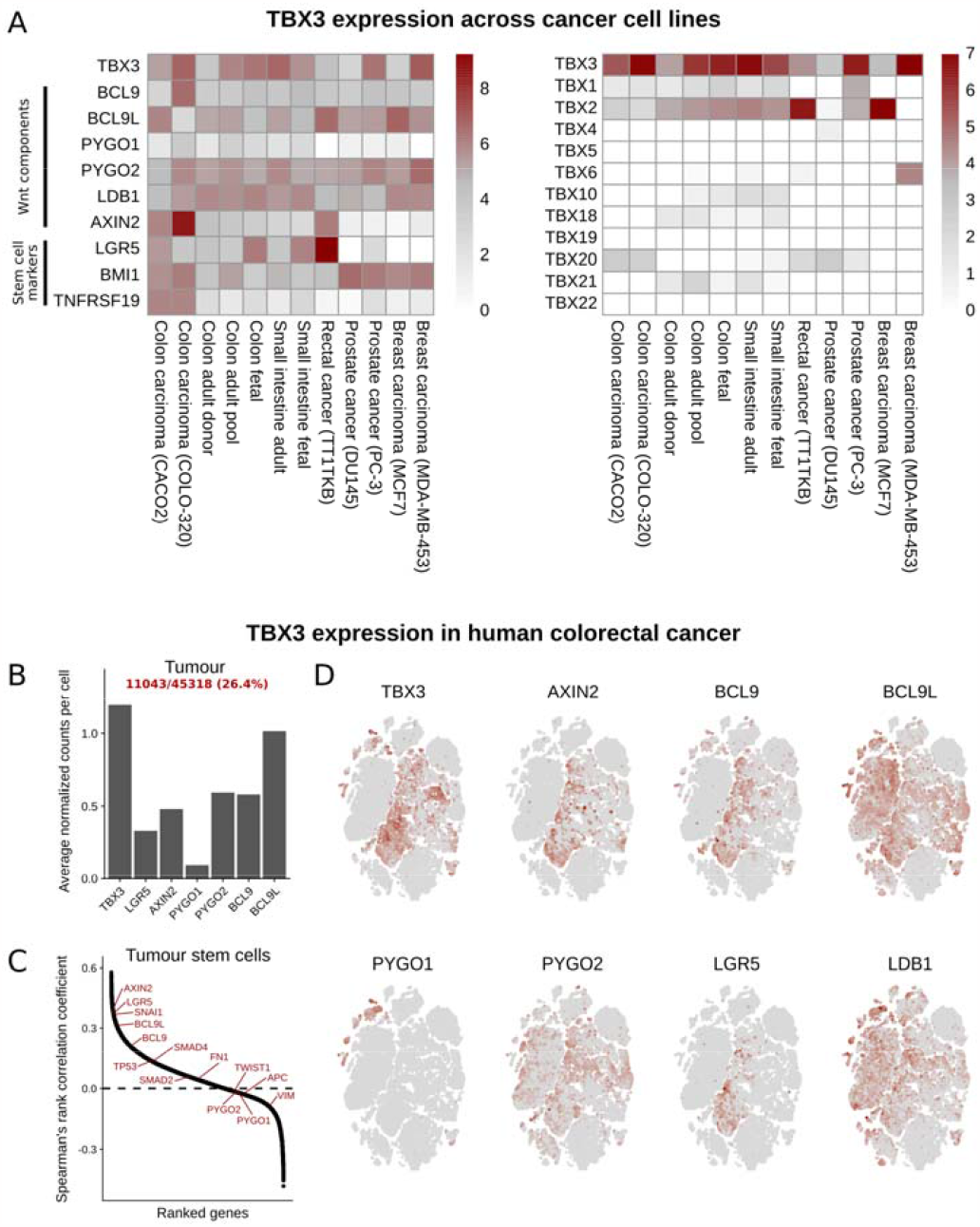
TBX3 expression in human colorectal cancer. (A) Heatmaps showing normalized expression of TBX3 for a set of cancer cell lines. Left panel: expression of TBX3 and selecte Wnt-relevant genes of interest. Right panel: expression of all T-box transcription factors. CAGE-seq data generated by the FANTOM5 project, and datasets was retrieved through the FANTOM5 Table Extraction Tool (https://fantom.gsc.riken.jp/5/tet/#!/search/hg19.cage_peak_ann.txt.gz). (B) Barplot showing average normalized counts detected in tumor tissue cells for TBX3 and the selected Wnt-relevant genes LGR5, AXIN2, PYGO1, PYGO2, BCL9 and BCL9L. (C) Ranked distribution of calculated Sp arman’s rank correlation coefficients between TBX3 and each other gene, based on cells identified a stem cells in tumor. The location of genes of interest are annotated with red colored text in the plot. () t-distributed stochastic neighbor embedding (t-SNE) plots of tumor cells highlighting the expression pattern across cells. scRNA-seq data from Uhlitz and colleagues^35^

Single cell RNA sequencing (scRNAseq) of human CRC confirmed TBX3 expression in tumor cell populations corresponding to a considerable 26.4% of all tumor cells ^35^ (Figure 1B). Average normalized counts for *TBX3* correlated with counts of i) relevant Wnt targets (*AXIN2* and *LGR5*) that are also markers of intestinal stem cells, ii) genes encoding for Wnt/β-catenin signaling components (e.g., *PYGO1, PYGO2, BCL9* and *BCL9L*) and mesenchymal markers associated to metastasis (e.g., *SMAD4, FN1, TWIST*) (Figure 1C). Across tumor cell populations, *TBX3* expression significantly overlapped with that of *BCL9* and *BCL9L* in individual cells (Figure 1D). Notable was also the expression of *TBX3* in *LGR5*^+^ single cell compartment (Figure 1D), indicating that TBX3 is likely to function in intestinal epithelial stem cells or cancer stem cells originating from this tissue ^36^.

### TBX3 displays genome-wide physical association with the chromatin in human colorectal cancer cells

We set out to determine the activity of TBX3 as regulator of gene expression by examining its genome-wide binding pattern in the human CRC cell line HCT116, in which a gain-of-function mutation in *CTNNB1*, the gene encoding for β-catenin, causes constitutive activation of Wnt signaling. We employed CUT&RUN (C&R) with Low-Volume and Urea (LoV-U; Zambanini et al., 2022) and performed 25 experimental replicates to define via the ICEBERG approach the full set of TBX3 binding sites in human CRC cells ^38^ (Figure 2A). When individually inspected, all 25 C&R-LoV-U experiments appeared successful, as they invariably displayed high-signal peaks in previously identified TBX3 target genes (e.g. WNT9A, Zimmerli et al., 2020). Importantly, all replicates pointed to the identification of previously unknown TBX3 bound regions, including the relevant proto-oncogene *MET*, whose mutations have been previously associated to aggressive forms of familial CRC ^39^ (Figure 2B).

**Figure 2.**
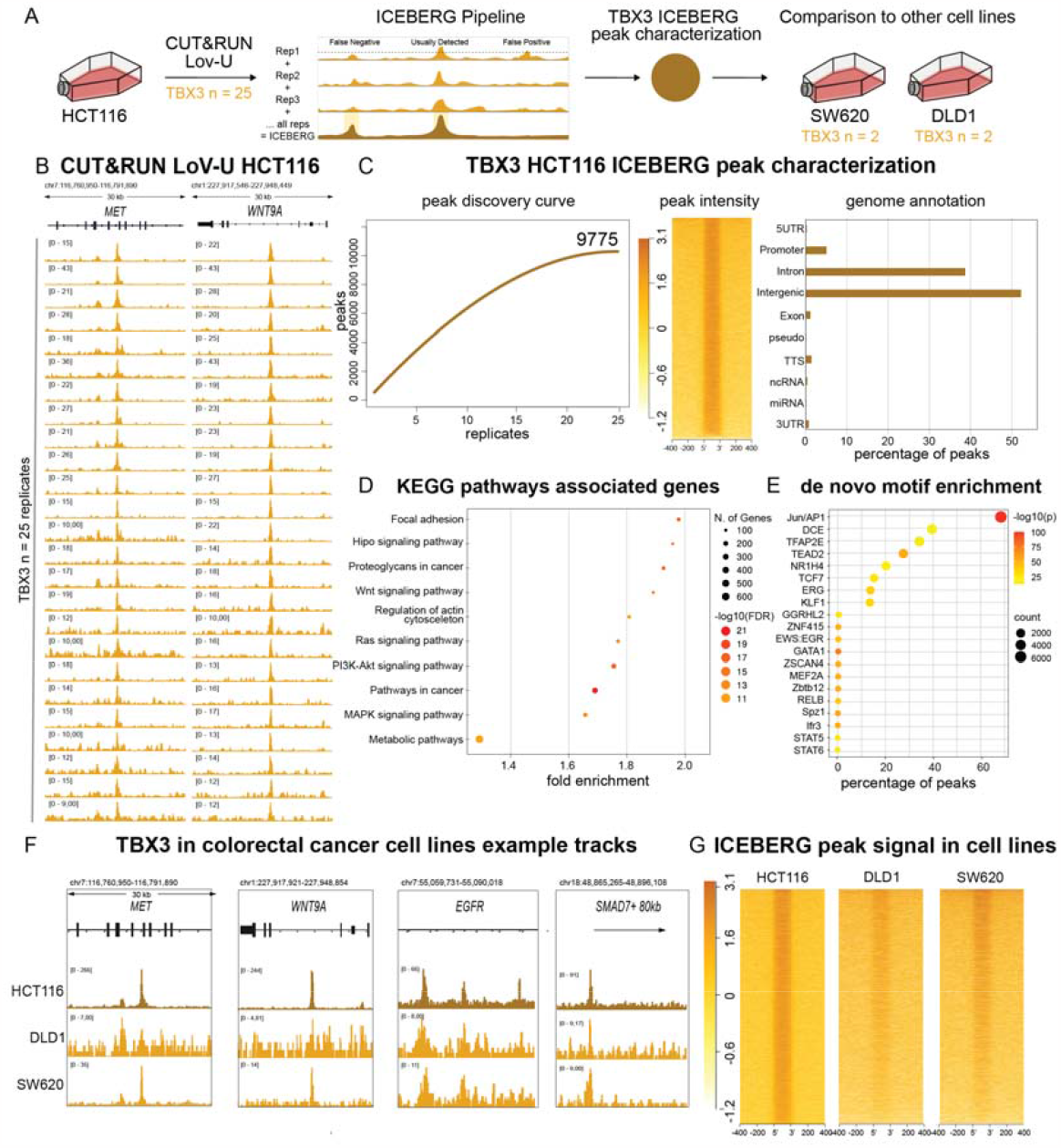
Genome-wide binding of TBX3 to the chromatin in colorectal cancer. (A) Schematic representation of the experimental design. 25 replicates of CUT&RUN LoV-U were performed against TBX3 in HCT116 human colorectal cancer cells. The ICEBERG pipeline was used to process the data, which was then compared to two other CRC cell lines: SW620 and DLD1. (B) Genome browser tracks showing high signal to noise ratio for all 25 TBX3 replicates at the MET and WNT9A loci. (C) Right: polynomial regression curve of the rate of peak discovery in the TBX3 ICEBERG, which plateaus as it reaches saturation between 20 and 25 replicates. Middle: signal intensity plot showing the signal to noise ratio in the full set of TBX3 ICEBERG peaks. Right: genome annotation as performed by HOMER, indicating that most TBX3 peaks are in intergenic or intronic regions. (D) KEGG pathway enrichment of the set of TBX3 peak associated genes (gene annotation done with GREAT), showing enrichment for many cancer-related signaling pathways, including Wnt and MAPK. (E) De novo motifs discovered by HOMER in the TBX3 peaks. The TBX3 peaks are statistically devoid of enrichment for any T-box motif. (F) Signal intensity plots of TBX3 CUT&RUN signal within ICEBERG peaks, comparing HCT116 with DLD1 and SW620 cells. SW620 more closely recapitulates the signal of HCT116, though DLD1 also shows signal enrichment in most regions. (G) HCT116, DLD1, and SW620 TBX3 CUT&RUN genome browser tracks at the example loci MET, WNT9A, EGFR and SMAD7. CRC = colorectal cancer.

Our approach yielded a total of 9775 high-confidence peaks, with peak discovery rate decreasing between replicates 20 and 25 (Figure 2C, left), suggesting saturation of new finding and implying that our strategy is close to identify the complete set of TBX3 binding events in HCT116 cells ^38^. These sites were characterized by high signal-to-noise ratio (Figure 2C, middle panel), and were mostly found in intergenic regions (∼38%) and introns (∼53%), while a smaller fraction (∼5%) was located at promoter regions (Figure 2C, right panel).

The fact that TBX3 displayed such a broad genome binding activity in CRC cells was indicative of a functional contribution to the biological features of these cells. To investigate which features these might be we used GREAT to annotate peaks with one or more genes, depending on proximity or previously described regulatory elements ^40^. TBX3 peaks were annotated to 7575 genes largely associated with the Kyoto Encyclopedia of Genes and Genomes (KEGG) pathways that describe cell mobility and malignant progression, including Wnt signaling, regulation of actin cytoskeleton and the Ras pathway, among others (Figure 2C, bottom left). This corroborated our previous identification of TBX3 as functional enhancer of metastatic dissemination ^26^.

Motif analysis of the DNA sequences underlying the TBX3 binding sites revealed a plethora of transcription factor signatures, including those for the TCF/LEF, TEAD and Jun/AP1 regulators (Figure 2E). Surprisingly, we could not identify statistical enrichment for TBX motifs, suggesting that, in this context, TBX3 likely plays a role as a cofactor hijacking preassembled transcriptional complexes. When C&R-LoV-U against TBX3 was performed in other human CRC cells, such as DLD1 and SW620 that, instead of activating mutations in *CTNNB1* carry loss of function alleles of *APC* ^41^, comparable enrichment for TBX3 signal was observed, both in the proximity of known TBX3 target genes (Figure 2F) as well as genome-wide (Figure 2G). As these CRC cell lines have been shown to be representative models of the main molecular subtypes or primary CRC, both concerning their mutational profile and their gene expression ^42^, our ICEBERG dataset of TBX3 target regions in different CRC cell types likely represents the general genomic activity of TBX3 in human CRC.

### The TBX3 binding profile largely overlaps with that of the Wnt signaling mediator β-catenin

From these and previous results obtained in the embryonic murine forelimb ^26^, we hypothesized that TBX3 could cooperate with β-catenin also in CRC cells. This hypothesis would imply that the TBX3 bound regions in CRC cells (Figure 2) would match, to a certain extent, β-catenin occupancy genome-wide in the same cells. To test this, we measured the peak overlap identified by the TBX3 ICEBERG with those found by the β-catenin ICEBERG (also 25 replicates in HCT116, Nordin et al., 2023) to identify all the regions bound both by TBX3 and β-catenin. Indeed, 53% of the identified TBX3 binding sites (5176 out of 9775 peaks) were also bound by β-catenin, indicating that more than half of TBX3 genome-wide activity is in concert with the Wnt/β-catenin transcriptional complex (Figure 3A). Importantly, even the TBX3-only and the β-catenin-only peaks displayed minor signal enrichment for the other factor – β-catenin or TBX3, respectively (Figure 3B), suggesting that our stringent peak calling parameters and overlap definition (at least one nucleotide in common) might underestimate an even stronger interplay between TBX3 and β-catenin. The highconfidence TBX3-β-catenin common peaks (Figure 3B, center) exhibit the strongest signal when compared to those that are, according to peak calling, exclusive to one or the other factor (Figure 3C).

**Figure 3.**
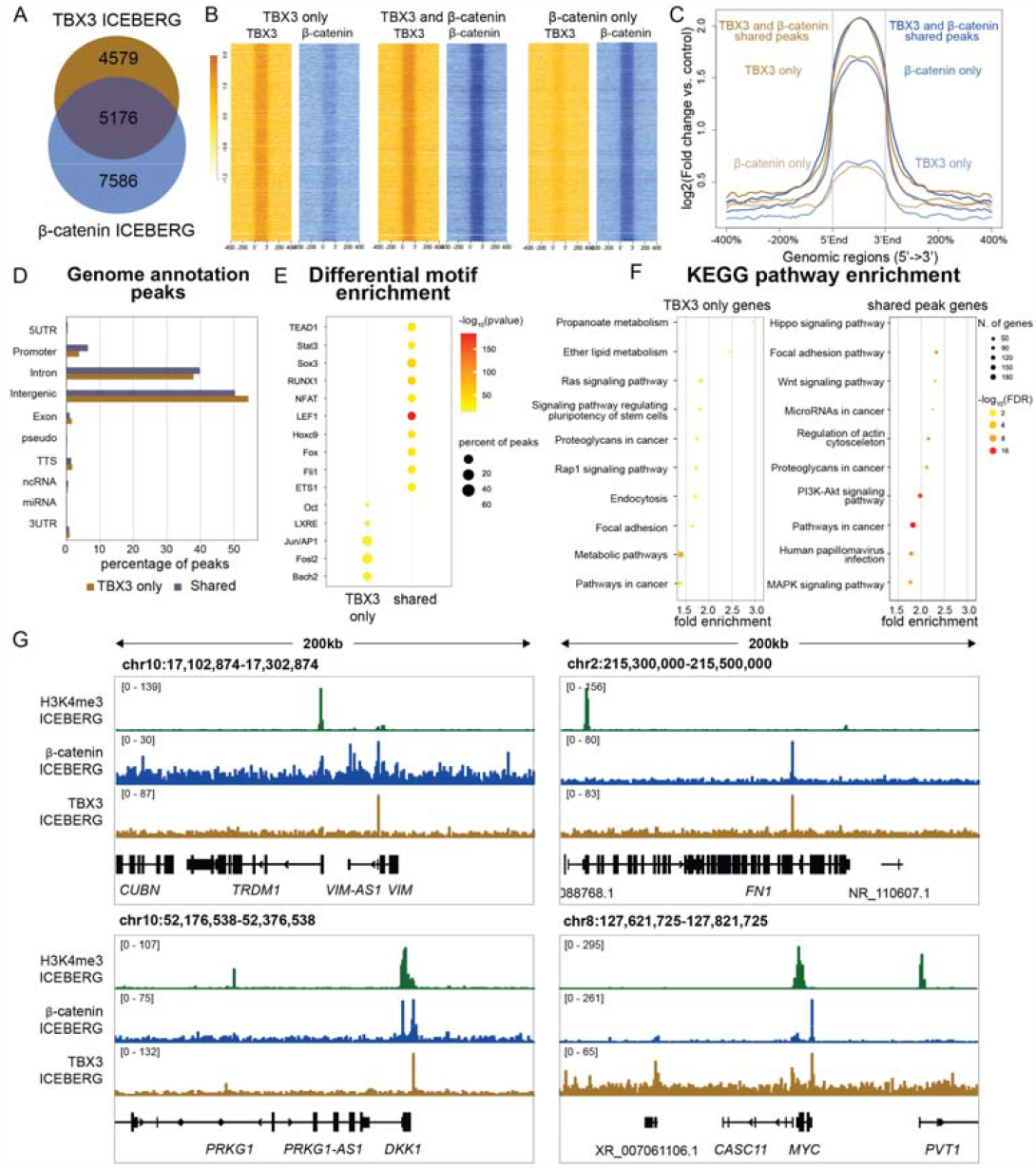
TBX3 and b-catenin have a partially overlapping binding profile in colorectal cancer. (A) Venn diagram depicting the overlap between ICEBERG peak regions of TBX3 and β-catenin in HCT116 cells. Over 50% of TBX3 peaks are also bound by β-catenin. (B) Signal intensity plots of TBX3 and β-catenin CUT&RUN signal in TBX3 only peaks (left), shared peaks (center) and β-catenin only peaks (right). Both TBX3 and β-catenin exhibit the strongest signals within the peaks that they share, compared to peaks exclusive to one or the other. (C) Signal profiles of TBX3 and β-catenin CUT&RUN signal in TBX3 only peaks, shared peaks, and β-catenin only peaks. (D) Genome annotations by HOMER showing that there are not large differences between the locations of TBX3 only peaks versus those shared with β-catenin. (E) Differential motif enrichment done with HOMER between TBX3 only peaks and shared peaks. TBX3 only peaks are differentially enriched for Jun/AP1 family factors, while shared peaks are highly significantly enriched for the Wnt signaling transcription factor LEF1, as well as others. Neither set individually shows statistical enrichment for Tbox motifs. (F) KEGG pathway enrichment for peak associated gene sets, comparing TBX3 only to TBX3 and β-catenin shared peaks. Shared peaks are enriched for the Wnt and Hippo pathways, while TBX3 only peaks are enriched for Ras. Both have enrichment for general pathways in cancer, and for focal adhesion. (G) Visualization of the relevant target loci displaying β-catenin and TBX3 CUT&RUN overlapping signal (blue and brown tracks, respectively). Green tracks show enrichment for H3K4me3, marking active promoter regions of the highlighted *VIM, FN1, DKK1* and *MYC* genes.

Genomic annotation of the TBX3-bound regions by HOMER identified prominent association with intergenic and intronic regions and found no obvious differences between the locations where TBX3 binds alone versus those shared with β-catenin (Figure 3D). TBX3-only peaks are enriched for Jun/AP1 family factors, among other, while shared TBX3 and β-catenin peaks presented enrichment for the Wnt signaling transcription factor LEF1 (Figure 3E). The absence of T-box motifs appearing in this analysis, in conjunction with the enrichment of the TCF/LEF signature, suggested that also in CRC cells, as previously found in developing mouse forelimbs ^26^, the association of TBX3 to these loci depends on the physical interaction of TBX3 with components of the Wnt/β-catenin transcriptional complex. KEGG pathway analysis of genes associated to peaks that present both TBX3 and β-catenin binding showed enrichment for biological processes related to regulation of cell migration and proliferation, including remodeling of actin cytoskeleton, focal adhesions, and the Hippo, Ras and Wnt signaling pathways (Figure 3F), raising the possibility that TBX3 is involved in the direct regulation of these groups of genes in CRC. Notable common TBX3/β-catenin targets included the epithelial-to-mesenchymal (EMT) genes *VIM* and *FN1*, both recognized drivers of malignant transformation of CRC ^43,44^, the general Wnt target *DKK1*^45^, and the CRC-driver proto-oncogene *MYC* ^46^ (Figure 3G).

#### TBX3 regulates genes that promote CRC cell metastatic behavior

The genome-wide binding profile of TBX3 indicated that this protein might directly regulate genes involved in metastatic behavior directly (Figures 2-3). This is important, as it could explain our previous observation that TBX3 overexpression in HCT116, which carry a constitutively active β-catenin, was sufficient to enhance metastatic cell dissemination in *in vivo* models ^26^. Moreover, others have identified that overexpression of TBX3 in CRC correlated with poor prognosis of CRC ^47^. To discover the relationship between TBX3 physical chromatin-binding and the observed enhancement of metastatic cell behavior, we set out to measure which genes are transcriptionally controlled by TBX3 in CRC. We employed Cap-Analysis Gene Expression sequencing (CAGE-seq), which measures the abundance of RNAs and maps the transcriptional start sites (TSS) and other 5’-capped regulatory regions, that include putative regulatory elements ^31,48^ (Figure 4A). CAGE-seq revealed a total of 155 differentially expressed transcripts in HCT116 upon TBX3 overexpression compared to control empty vector condition (Figure 4B): of these, 129 were upregulated and 26 downregulated, possibly indicating that, in this context, TBX3 primarily mediates gene activation rather than gene repression ^49^.

**Figure 4.**
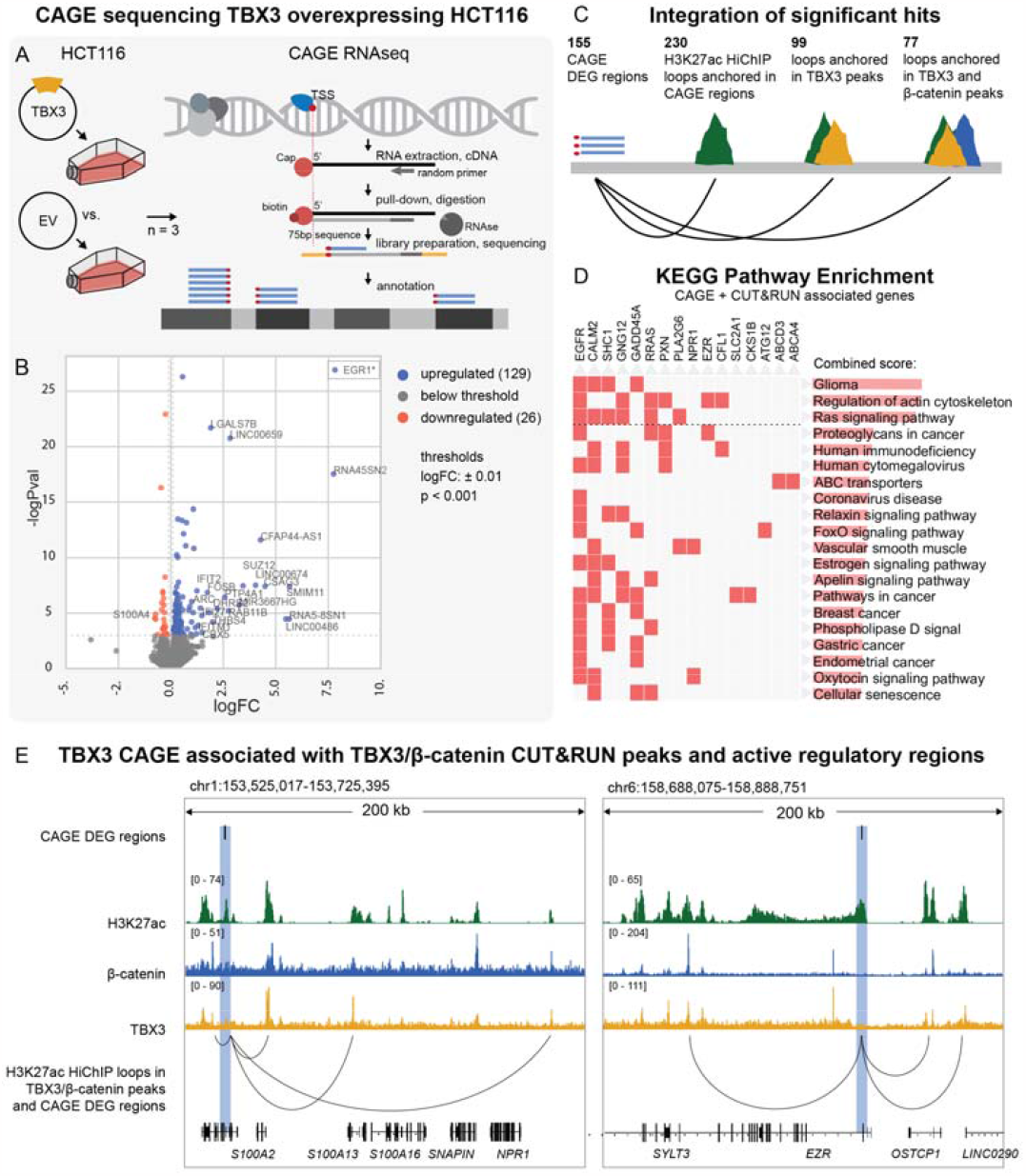
CAGE-seq identifies TBX3-regulated drivers of metastasis. (A) Schematic representation of the experimental setup, where TBX3 was overexpressed in HCT116 cells, and the performed procedure for Cap-Analysis Gene Expression Sequencing (CAGE-Seq). CAGE-seq was used to identify both differentially expressed genes, their exact transcription start sites, and regions of transcribed enhancer RNAs. (B) Volcano plot of CAGE-seq differentially expressed genes (DEG) regions (upregulated in blue (p<0.001, logFC>0.1), down regulated in red (p<0.001, logFC<-0.1), excluding TBX3 itself. 155 regions were found enriched. EGR1 transcript (-logP= 89.04, logFC=2.1) was visualized separately to increase visibility. (C) Schematic of the strategy used to identify genomic elements regulated by TBX3 and TBX3b-catenin. H3K27ac-HiChIP data identified DNA-DNA connections between active regulatory regions, including promoters and putative enhancers. We focused on the differentially expressed RNAs (CAGEseq) that emerge from regions of the genome that are connected via H3K27ac-marked regions (green) to TBX3 (orange) and/or b-catenin (blue) CUT&RUN peaks. (D) KEGG pathway enrichment of the genes annotated to CAGE-seq DEG loops, showing high fold enrichment for glioma, regulation of actin cytoskeleton, and the Ras pathway. Bar graphs visualize a combined score (log(p)*z-score). Dotted line illustrates the threshold of p< 0.05. (E) Example genome browser visualization of CAGE-seq DEG regions that are connected via H3K27ac mediated chromatin loops to TBX3 and b-catenin shared CUT&RUN peaks, in cancer relevant loci near S100 genes and EZR. DEG = differentially expressed gene.

We set out to understand which of these changes in RNA abundance could be directly controlled by TBX3. To this aim, we integrated the TBX3 genomic distribution with the network of active CRC-enhancers obtained by HiChIP targeting the marker of open chromatin H3K27ac in HCT116 ^50^. The 155 differentially expressed CAGE-seq regions were anchored to 230 H3K27ac-mediated DNA-DNA interaction loops (Figure 4C; note that each CAGE-seq region could be anchored to >1 H3K27ac loop), supporting the notion that they correspond to active regulatory elements ^51^. Among the many H3K27ac-ornamented regions looping into differentially expressed CAGE-seq RNAs, a large fraction (99 out of 230, 43%) was occupied by TBX3, and the majority of these (77 out of 99, 78%) was co-occupied by both TBX3 and β-catenin (Figure 4C), indicating that TBX3 and β-catenin together promote the differential RNA production measured by CAGE-seq.

KEGG pathway enrichment analysis of the TBX3/β-catenin regulated genes showed high-fold enrichment for cancer related processes, including regulation of actin cytoskeleton and the Ras pathway (Figure 4D). Integration of CAGE-seq, TBX3 C&R-LoV-U, β-catenin C&R-LoV-U, and HiChIP targeting H3K27ac permitted us to identify the specific and important instances of direct gene regulation by TBX3 and β-catenin in CRC. These include cancer relevant loci near the *S100* family of genes, which are considered prognostic markers for patients with colorectal neoplasia ^52^, and *EZR*, a known mediator of invasion and metastasis ^53,54^ (Figure 4E).

### TBX3 associates with the Wnt/β-catenin transcriptional complex

One observation that attracted our attention was that TBX3 seemed to associate to the chromatin prominently, but not via its DNA-binding motif. We reasoned that TBX3, in CRC, might act via hijacking other chromatin-associated protein complexes. To identify its molecular mechanism of action in CRC, we set out to screen for TBX3-interacting nuclear proteins. We selected the Wnt-responsive HEK293T cells, which allowed us to modulate Wnt/β-catenin signaling either by the GSK3 inhibitor CHIR99021 (CHIR) to activate the pathway (WNTON) ^55^ or by the Porcupine inhibitor LGK974 (LGK) to inhibit the endogenous secretion of WNT ligands (WNT-OFF) ^56^, and employed the BioID technology ^57^ to map the physical proximities of TBX3 in both conditions (Figure 5A, Figure S1A). TBX3 was cloned in frame to the biotin-ligase BirA and expressed in HEK293T cells cultured in WNTON and WNT-OFF conditions. Biotinylated nuclear extracts were subjected to streptavidin pulldown, and nuclear proteins in the vicinity of TBX3 before and after Wnt activation were isolated and identified via mass spectrometry. A total of 517 candidate interactors were found in CHIR-treated cells and 78 candidate interactors were found in LGK-treated cells (SAINT score > 0.6, BFDR < 0.05) (Figure 5A). Among the proteins that were the most highly enriched in CHIR-treated cells we identified several known regulators of gene expression associated to Wnt signaling, such as ARID1B, SMARCA1 and SMARCE1, JUN/FOS, MED13, as well as the member of the TCF/LEF family of transcription factors TCF7 (Figure 5B). Activation of Wnt signaling by CHIR seemed to enhance the vicinity of TBX3 to all these proteins (Figure 5B, right panel), possibly reflecting the broad stabilization of the proteome that follows GSK3 inhibition ^58^. In WNT-ON conditions, both Gene Ontology (GO) and KEGG identify the TBX3 vicinal proteins as being enriched in positive regulation of transcription, chromatin remodeling, and Wnt signaling pathway, among other terms (Figure 5C). CORUM analysis of protein complexes ^59^ identified the prominent association of TBX3 with the SWI/SNF (BAF) chromatin remodeler ^16^, the Mediator ^60^ and LDB1 (LIM domain-binding protein, ^15^), all known to assist the transduction of Wnt target genes (Figure 5C, right panel). These experiments support a model in which TBX3 is tethered to the WREs by direct and promiscuous physical contacts with members of the Wnt/β-catenin transcriptional complex during Wnt pathway activation.

**Figure 5.**
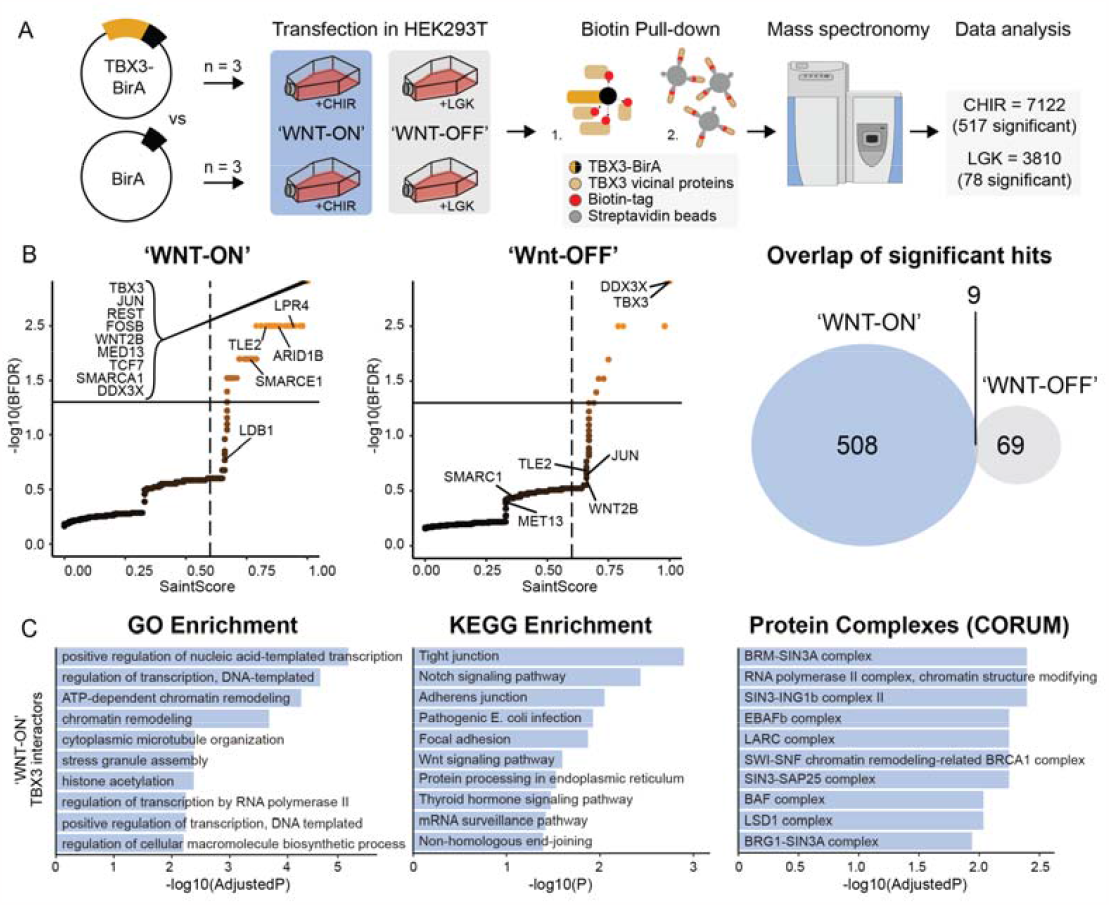
BioID reveals the effect of WNT activation on TBX3 protein-proximity network. (A) Schematic representations of the experimental design. HEK293 cells expressing a TBX3-fused biotin ligase (BirA) versus an empty vector-BirA, were cultured in either CHIR (‘WNT-ON’) or LGK (‘WNT-OFF’) supplemented media (n=3). Biotinylated proteins were pulled down and further subjected to mass spectrometry analysis. (B) left panel: Scatter plots displaying the mass spectrometry results of the ‘WNT-ON’ and ‘WNT-OFFf’ conditions (thresholds are SaintScore > 0.6, BFDR < 0.05). right panel: Venn diagram indicating common (9) and unique (508 for ‘WNT-ON’, 69 for ‘WNT-OFF’) TBX3-proximity labeled proteins in both conditions. (C) Enrichment of ‘WNT-ON’ unique vicinal proteins of TBX3 in Gene Ontology terms (GO), Kyoto Encyclopedia of Genes and Genome pathways (KEGG), and protein complexes (Comprehensive Resource of Mammalian protein complexes, CORUM).

### A conserved NPF motif mediates TBX3 interactions to the Wnt/β-catenin transcriptional complex

Investigation of the amino acid residues composing TBX3 revealed an NPF (Asparagine-Proline-Phenylalanine) motif adjacent to the DNA binding T-box domain (Figure 6A, S2A). The NPF caught our attention for three reasons. First, this NPF motif was among the most conserved sequences of TBX3 and across the members of the TBX family of transcription factors (Figure 6A, S2B; ^61^). Second, an NPF motif was first described as the transactivation domain of the Wnt co-factor Pygopus (PYGO1 and PYGO2 in human) ^62^ and recent structural work revealed that PYGO-NPF mediates its interaction with the chip/LDB-SSDB (ChILS) complex in the Wnt/β-catenin transcriptional complex ^63,64^. Third, our original identification of TBX3 as facultative member of the Wnt/β-catenin transcriptional complex derived from genetic data implying TBX3 function in PYGO-independent phenotypes ^26^.

**Figure 6.**
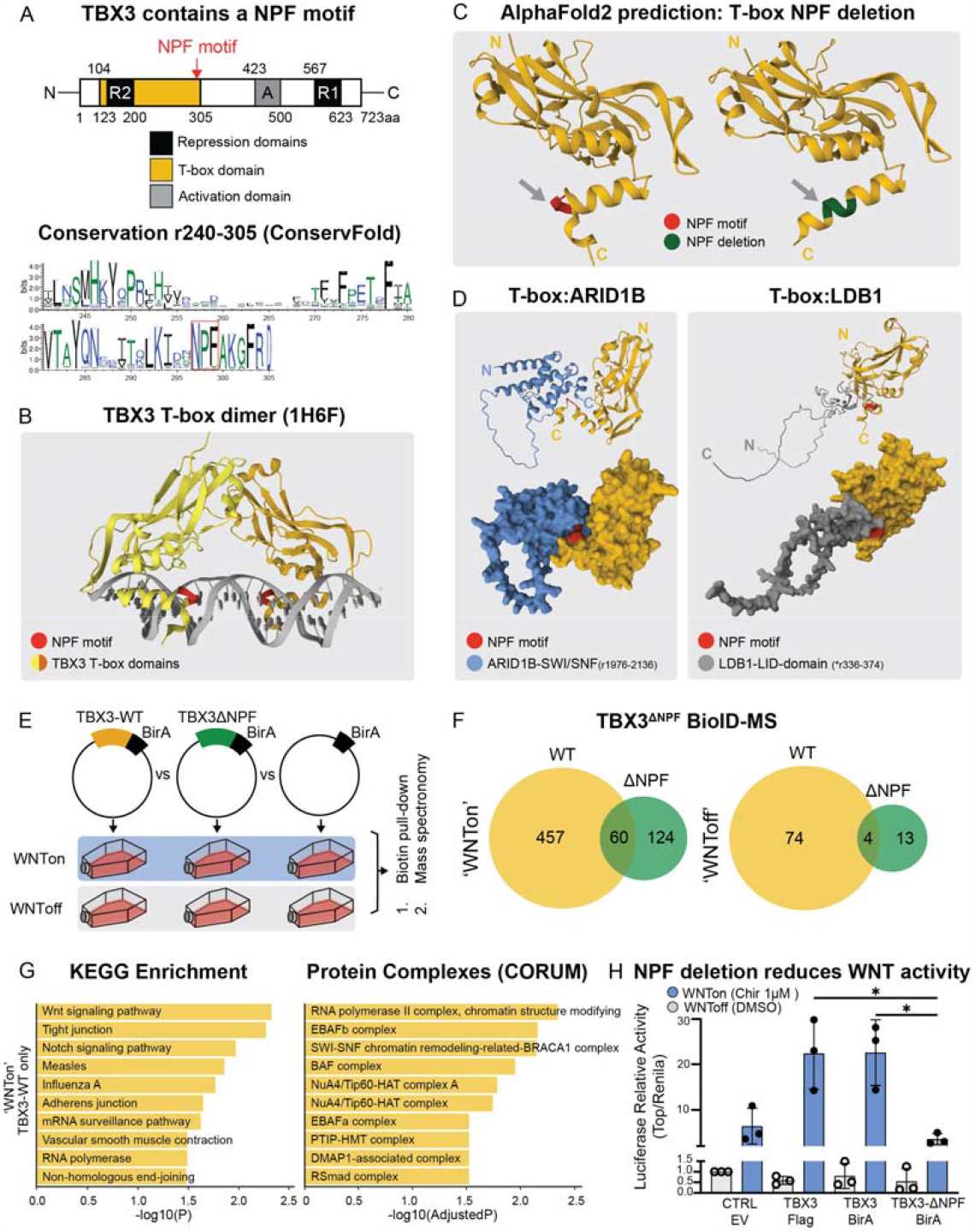
TBX3’s NPF-motif engages in protein-protein interaction unique to ‘WNTon’ condition. (A) TBX3 features a conserved NPF-motif, as indicated by the WebLogo conservation sequence predicted by ConservFold (lower panel, residue 240-305). The y-axis indicates information content in bits, with the overall height of the characters based on positional conservation. The color scheme represents the hydrophobicity of the amino acids, with hydrophilic in blue (RKDENQ), neutral in green (SGHTAP), and hydrophobic in black (YVMCLFIW). The NPF is located at the Cterminal end of its T-box domain. (B) 3D structure of the experimentally determined, DNA-bound T-Box domain (PDB 1H6F) dimer of human TBX3 61 with the NPF-motif highlighted in red. (C) AlphaFold2 prediction of the T-Box domain of human TBX3 with and without deletion of the NPF motif. The NPF-motif in red, area spanning the deletion in green. (D) AlphaFold2 predictions of interaction of two exemplary proteins from TBX3 vicinal protein that were unique to the ‘WNT-ON’ condition (from Figure 5). Left: Predicted interaction between TBX3 T-box domain (yellow) and ARID1B SWI/SNF domain (gray). Right: Predicted interaction between TBX3 T-box domain (yellow) and LDB1 LID-domain (gray). NPF-motif is shown in red. (E) Schematic illustration of the experiment investigating changes in TBX3 vicinal proteins upon deletion of its NPF motif (TBX3-DNPF). HEK293T cells were transfected with TBX-BirA, TBX3-DNPF-BirA and empty vector, respectively, in ‘WNT-ON’ and ‘WNT-OFF’ conditions (n=3). Biotinylated proteins from the pull-down were further subjected to mass spectrometry analysis. (F) The number of common and unique vicinal proteins of TBX3 wild-type (TBX3-WT) and TBX3-DNPF in ‘WNT-ON’ (left) and ‘WNT-OFF’ (right) conditions, visualized as Venn diagrams (SaintScore > 0.6, BFDR < 0.05). (G) Enrichment of KEGG pathways and CORUM protein complexes in hits that are unique to the TBX3-WT vicinity in ‘WNT-ON’ condition only. (H) TOPFlash assay investigating the effect of TBX3 NPF deletion on the activation of Wnt/b-catenin signaling. HEK293T cells were transfected either with control empty vector (EV), or TBX3-Flag, or TBX3-BirA or TBX3-BirA-DNPF. All cells were transfected with the TCF/LEF-luciferase reporter SUPERTOPflash and Renilla control plasmids. WNT-OFF (gray) and WNT-ON (blue) conditions were tested. Activation of Wnt signaling was measured as relative luciferase activity. SEM. *p<0.05.

Structural data of the dimeric TBX3 DNA-binding domain reveals that the NPF is an exposed “anchor” between two alpha-helices at the C-terminus of the T-box domain (Figure 6B, S2B) ^61^. AlphaFold2 (AF2) structure modeling predicts perturbation of the local 3D structure upon deletion of the NPF (Figure 6C), supporting the notion of this motif being a relevant stretch of the protein whose deletion however does not perturb general folding. As our genome-wide binding profile hints at TBX3 association to the chromatin via protein-protein interaction, instead of direct contact to the DNA, we hypothesized that the TBX3-NPF, as it occurs for PYGO, could mediate interactions with other proteins. AF2 Multimer modeled interactions with several components of the BAF and ChiLS complexes, including ARID1B and the LID domain of LDB1, where the NPF lay adjacently to the interaction surfaces (Figure 6D). According to AF2 Multimer, the dimeric DNAbinding domains of TBX3 could be positioned in a complex which includes LDB1, SSBP2 and the HD1 domain of BCL9; in this model, the two TBX3-NPF motifs are close to LDB1 and surround a groove that could, in our opinion, host the DNA and other DNA-binding proteins, such as ARID1B (Figure S2C, D).

To validate our prediction ^65^, we set out to test the consequence of deleting the NPF from TBX3 on its ability to interact with its protein partners. We cloned a mutant version of TBX3 only lacking these three NPF amino acid residues (TBX3ΔNPF) in frame with the BirA biotinylating enzyme and carried out a BioID assay similar to the one described above. Both in WNT-ON as well as WNT-OFF conditions, TBX3ΔNPF lead to a dramatically decreased identification of significant peptide hits, indicating a reduced affinity with proteins partners (Figure 6F). As the BioID was done on the nuclear fraction, this indicates that the TBX3ΔNPF variant did not lose its nuclear translocation capacity. Among the lost interactors were the components of the BAF and ChiLS complexes, including ARIDB1 and LDB1, supporting the hypothesis that their interaction with TBX3 is mediated by the NPF motif of TBX3 (Figure 6G, S2E).

While deletion of the NPF motif affected the interaction with these relevant proteins, the exposure of this motif at the C-terminus of the T-box domain and its proximity to both protein partners and the DNA determined by both crystal (Figure 6B; Coll et al., 2002) and AlphaFold2 model (Figure 6D, S2C) structures, left the possibility open that the deletion of NPF affected not only protein-protein interactions, but also its association with the DNA on the TBX consensus sequence. To distinguish between these possibilities, we employed the transcriptional superTOPFLASH reporter that specifically and exclusively responds to TCF/LEF mediated transcription upon Wnt pathway activation ^66^. We have previously shown that TBX3 could amplify the Wnt-dependent transcription at suboptimal stimulation (1 μM) with CHIR ^26^. Here we tested the consequence of NPF deletion in this assay. TBX3-WT-BirA, similarly to the TBX3-FLAG, significantly enhanced transcription from the Wnt reporter superTOPFLASH (Figure 6H), showing that the BirA moiety does not perturb TBX3 activity or folding. TBX3ΔNPF-BirA, on the other hand, had no effect on superTOPFLASH transcription (Figure 6H), showing that TBX3 effect on the Wnt-dependent transcription is mediated by the NPF and its ability to confer interaction with the components of the Wnt/β-catenin transcriptional complex.

## Discussion

Revealing the molecular composition of the transcriptional complexes that drive gene expression is a primary objective to understand how cell behavior is regulated during embryonic development, cell homeostasis, and how disease results when such mechanisms are perturbed. Our primary objective is to understand how the Wnt signaling pathway, with its apparent universal transduction mechanism, can drive diverse gene expression patterns across a wide range of developmental and homeostatic contexts ^2,67^. This, we posit, will be key to unravelling what are the relevant cell-specific transducers that confer oncogenic potential to the Wnt/β-catenin axis ^68^. Addressing this knowledge gap is pivotal since neoplasia of the colorectal epithelium begins with aberrant Wnt/β-catenin signaling and broadly targeting the Wnt pathway can lead to undesirable consequences.

We have previously discovered that the tissue-specific and human disease-relevant transcription factor TBX3 can assist the BCL9/β-catenin driven transcriptional complex downstream of Wnt signals during mouse forelimb development ^26^. Our finding opened the possibility that TBX3 is a relevant player in other tissues, and that the physical interplay with the Wnt pathway (Figure 7) might underlie the oncogenic activity that had been associated to its aberrant expression in CRC and other cancers ^34^. Our study provides a primary example: we find that TBX3 is expressed in subsets of CRC cells, likely as a consequence of hyperactive Wnt signaling, as its expression overlaps with that of other Wnt target genes and Wnt signaling components, as well as markers of the intestinal stem cells. Moreover, we detected TBX3 engaging with transcriptional complexes positioned genome-wide at regulatory sites along the chromatin, modulating the expression of genes that are known to drive CRC metastatic dissemination. The identification of the molecular activities of TBX3 in human CRC cells not only explains how its overexpression can increase their metastatic potential, as we previously observed, but also establishes TBX3 as a new regulator of metastasis and as a putative target for molecular therapy.

**Figure 7.**
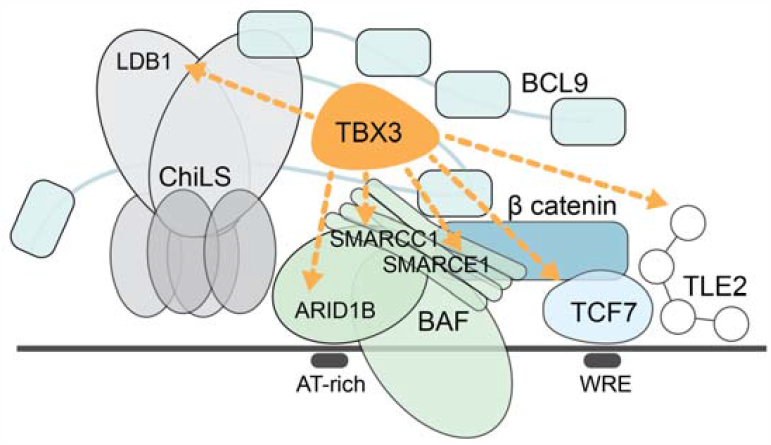
Schematic model of TBX3 interaction with the Wnt enhanceosome. TBX3 interacts with several components of the Wnt enhanceosome. TCF7 and TLE2 are nuclear components of Wnt signaling (blue). ARID1B, SMARCC1 and SMARCE are components of the BAF complex (green). LDB1 is part of the ChiLS complex. WRE = Wnt responsive elements.

Identification of the NPF motif, at the C-terminal end of the DNA-binding T-box domain, further supports the impetus for considering TBX3 as a potential molecular target. While more than 30 years of evidence have supported a critical role of abnormal Wnt/β-catenin activation as causative of CRC ^69,70^, the field is still missing Wnt pathway inhibitors for therapeutic treatment of this type of cancer ^5–8,71^. The NPF motif appears to be present in several other proteins and have a defined mechanism of action ^72^. Many of these proteins, such as Pygopus and ARIDB1, possess oncogenic activity ^73^. Given its accessibility on the protein surface of TBX3 and its functional significance, the NPF of TBX3 could certainly constitute an attractive target for anti-cancer therapeutics. It remains to be established whether TBX3 is essential for intestinal stem cells homeostasis as β-catenin ^74^, or if its presence is dispensable as is the case for BCL9 and BCL9L ^75^. While we consider the latter hypothesis more plausible, since the phenotypes caused by loss of *Bcl9/9l or Tbx3* were similar in the mouse – an observation that drew us onto studying TBX3 in the first place – only functional or knockout *in vivo* experiments will provide a final answer. These will be required to assess whether TBX3 functions, in other biologically relevant contexts such as in pluripotent stem cells ^76^, is also mediated by its interplay with the protein complexes identified in the current study.

Intriguingly, inspection of published data on protein-protein interactions in colon cancer cell lines revealed that significant interactions between TBX3 and β-catenin had already been detected ^77^; we consider this an important independent validation of our study. The same authors, in a previous prominent article, had identified an interplay between β-catenin and another member of the TBX family of transcription factors, TBX5, which together with YAP1 and β-catenin was essential to mediate survival and tumorigenesis of several β-catenin-driven cancers, including CRC ^78^. Our failed attempts in introducing loss-of-function mutations in the *TBX3* gene (Zimmerli et al., 2020 and this study) support the notion that also TBX3, similarly to TBX5, might be essential for survival of CRC cells. This is important, as, if TBX3 proved to be dispensable for homeostasis as BCL9/9L but required for cancer cell survival as TBX5 or β-catenin, its targeting will constitute the proverbial therapeutic window.

## Acknowledgments

The authors are grateful to Dr. Hurlin for kindly donating the original TBX3 expressing plasmid, to Drs. Utpal P. Davé and Justin H. Layer for providing the LDB1 encoding plasmid. This work was supported by Grants to C.C. from Cancerfonden (CAN 2018/542 and 21 1572 Pj), the Swedish Research Council, Vetenskapsrådet (2021–03075), Linköping University and the Knut och Alice Wallenbergs Stiftelse. Grants from LiU-Cancer support collaborative efforts across the XF.S., E.vC. and C.C. laboratories. F.L., S.K., E.vC. and C.C. are Fellows of the Wallenberg Molecular Medicine (WCMM) and receive generous financial support from the Knut and Alice Wallenberg Foundation. A.J-M. was supported by grants from the Japan Society for the Promotion of Science (JSPS) in Japan, and the Capstone Awards from Stiftelsen för internationalisering av högre utbildning och forskning (STINT) in Sweden. The computations and data handling were enabled by resources provided by the National Supercomputer Centre (NSC), funded by Linköping University. Peter Münger at the National Supercomputer Centre is acknowledged for assistance concerning technical and implementational aspects in making the codes run on the Sigma resource.

## Author contributions

A.J-M., S.S. and C.C. conceived the work and designed the study. A.J-M., S.S., T.W., A.N., V.G., S.B., G.Z., J.U., A.M., E.F., L.M., performed the experiments. S.S., T.W., A.N., Y.vdG. and R.D. conducted the formal analyses. P.P., F.L., S.K., A.M. X.F.S., E.vC. and G.S. provided experimental and conceptual input. S.S., T.W. and A.N. designed the figures. C.C. supervised the research team, provided financial support, and wrote the manuscript with input from all authors.

## Competing interest statement

The authors declare no competing interests.

## Availability of Data and Materials

CAGE-seq raw and processed data have been deposited to ArrayExpress and can be accessed via accession number EMTAB-13647. CUT&RUN-LoV-U raw and processed data have been deposited to ArrayExpress and are available via the accession number E-MTAB-13646. Proteomics Data are available via ProteomeXchange with identifier PXD047899.

## Methods

### Cell culture

HCT116, DLD1, and SW620 human colorectal cancer cells, as well as HEK293T human embryonic kidney cells, were cultured in a 37 °C incubator in 5% CO2 and 89% humidity. Culture medium used was high glucose Dulbecco’s Modified Eagle Medium (Cat. #41965039, Gibco) supplemented with 10 % bovine calf serum (Cat. #1233C, Sigma-Aldrich) and 1X Penicillin-Streptomycin (Cat. #15140148, Gibco).

### CUT&RUN LoV-U

#### Protocol

CUT&RUN LoV-U was performed according to Zambanini et al., 2022, doing 25 replicates of TBX3 in HCT116 and 2 replicates each in DLD1 and SW620 cells, plus IgG negative controls. 250,000 cells/sample were harvested using Trypsin-EDTA (Cat. # 25200056, Gibco) for 5 - 10 minutes. Cells were washed in DPBS two times (Cat. #14190094, Thermo Fisher Scientific). Nuclear extraction was performed by three washes in Nuclear Extraction (NE) buffer (HEPES-KOH pH-8.2 [20 mM], KCl [10 mM], Spermidine [0.5 mM], IGEPAL [0.05%], Glycerol [20%], Roche Complete Protease Inhibitor EDTA-Free). After extraction, nuclei were resuspended in 40 µl NE per sample and bound to 10 µl Magnetic ConA Agarose beads equilibrated in binding buffer (HEPES pH 7.5 [20 mM], KCl [10 mM], CaCl2 [1 mM], MnCl2 [1 mM]) as described in Meers and colleagues ^79^. Bead binding proceeded for 15 min at 4 degrees, then beads were resuspended in 200 µl wash buffer per sample and distributed in PCR tubes. Samples were washed once in 200 µl EDTA wash buffer (wash buffer with EDTA [0.2 mM]), incubating 5 min at room temperature before being resuspended in antibody buffer (wash buffer with antibody 1:100). Incubation happened ON at 4 °C on a rotator. Antibodies used included anti-TBX3 ABIN6265491 and anti-rabbit ABIN101961. The next morning samples were washed 5 times and resuspended in 200 µl of pAG-MNase buffer (wash buffer with pAG-MNase 120ng/sample) and incubated for 45 min at 4 °C on a rotator. pAG/MNase was a gift from Steven Henikoff (Addgene plasmid #123461; http://n2t.net/addgene:123461; RRID: Addgene_123461) expressed and purified according to ^79^. Five washes were performed and during the last wash equilibrated 5 min in wet ice. Samples were resuspended in 200 µl ice cold wash buffer with 2 mM CaCl2for digestion of 30 min in wet ice. The digestion buffer was kept and transferred to tubes containing 1.5 µl of 0.5 M EDTA and 1.5 µl of 0.5 M EGTA to inactivate the pAG-MNase. Beads were resuspened in 47 µl of 1X Urea STOP buffer (NaCl [100 mM], EDTA [2 mM], EGTA [2 mM], IGEPAL [0.5%], Urea [8.8 M]) and the DNA was eluted for 1 hr at 4°C for elution. Beads were collected on the magnet and liquid containing DNA was transferred to the PCR tubes and mixed with the digestion buffer. DNA was purified using two successive rounds of bead purifications according to manufacturer’s directions, using MagBind TotalPure NGS beads (Cat. #M1327, Omega Bio-Tek) at 2X (200 µl first round and 40 µl second round), final elution was with 20 µl Tris-HCl pH 7.5.

#### Library Preparation and Sequencing

Library preparation was performed with the KAPA Hyper Prep Kit for Library preparation was performed with the KAPA Hyper Prep Kit for Illumina sequencing (Cat. #KK8504, KAPA Biosystems) according to manufacturer’s guidelines with modifications. 0.4X volume reactions were used for End repair and A-tailing steps. The thermocycler conditions were 12 °C for 15 min, 37 °C for 15 min and 58 °C for 25 min. Adapter ligation was also performed in 0.4X volume reactions. KAPA Dual Indexed adapters were used at 0.15 µM. DNA purification was performed after ligation with 1.2X volumes of Mag-Bind TotalPure NGS beads. Library amplification steps were performed in 0.5X volume reactions. The cycling conditions were set to: 45 sec initial denaturation at 98 °C, 15 sec denaturation at 98 °C, 10 sec annealing/elongation at 60 °C, 1 min final extension at 72 °C, hold at 4 °C. Libraries were amplified 13 cycles. DNA purification was performed with 1.2X beads. Libraries were size-selected using the E-Gel EX 2% agarose gel (Cat. # G402022, Invitrogen) and the E-Gel Power Snap Electrophoresis System (Invitrogen), selecting for fragments 150 - 500 bp. DNA from the gel was purified using the QIAquick Gel Extraction Kit (Cat. #28706X4, Qiagen) according to the manufacturer’s instructions. Libraries were quantified using Qubit (Thermo Scientific)’s high sensitivity DNA kit (Cat #Q32854, Thermo Scientific). Libraries were sequenced with 36 bp pair-end reads on the NextSeq 550 (Illumina) using the Illumina NextSeq 500/550 High Output Kit v2.5 (75 cycles) (Cat. #20024906, Illumina).

#### Data Processing and ICEBERG

Trimming was performed with bbmap bbduk (version 38.18) ^80^ to remove adapters, known artifacts, and repeats of AT, TA, or poly G/C. Alignment was done to hg38 with bowtie2 (version 2.4.5) ^81^, settings: – local –very-sensitive-local –no-unal –no-mixed -no-discordant –phred33 –dovetail -I 0 -X 500. SAMtools (version 1.11) ^82^ was used to fix improperly paired mates and for deduplication. Bam files were filtered with the CUT&RUN hg38 blacklist (suspect list) ^83^ using BEDTools (version 2.30.0) ^84^. BEDTools genomecov on pair-end mode was used to create bedgraphs, bedgraphs were visualized in IGV 85. Data was processed with the ICEBERG pipeline according to Nordin et al., 2023a, to generate the TBX3 ICEBERG dataset. Briefly, individual replicates were peak called with MACS2 (version 2.2.6) 86 with the options -f BAMPE --keep-dup all and -q 5e-2 against the control. The maximum number of usable fragments was determined by the smallest replicate, and each replicate was shuffled and downsampled to this depth 3X, then merged into aggregates. Each were separately peak called with MACS2 as described above. Between each replicate addition, peaks were called to generate the ICEBERG build curve, which was modeled by fitting polynomial regressions of increasing order (up to 5th) in R (version 4.2.2, lm function), selecting that which had the highest R2 value. Peaks called in at least 2 of the 3 aggregates were determined and filtered to remove peaks not called in at least one individual replicate at MACS2 -p 1e-2, to finally generate the final set of ICEBERG peaks for TBX3.

#### Downstream Analyses and Integration with Published Data

Signal intensity plots and profiles were generated using ngs.plot (version 2.63) 87. Genomic region annotation was done with HOMER (version 4.11) ^82^ annotatePeaks on default settings, and motif analysis was also done with HOMER with the -size given parameter. Annotation to genes was done using GREAT (version 4.0.4) 40 on default settings. Gene ontology and KEGG pathway analysis was done with ShinyGO 88. Intervene (version 0.6.4) 89 was used to create Venn diagrams.

CUT&RUN datasets of β-catenin were downloaded from Nordin et al., 2023a and overlapped with TBX3 ICEBERG peaks using bedtools or Intervene. HiChIP of H3K27ac in HCT116 data, performed by Chen and colleagues 50 was downloaded from GEO. Loops were filtered as described in Nordin et. al., and then filtered for those overlapping TBX3 peaks using BEDtools pairtobed. Bigwig files mapped to hg38 were downloaded for H3K27ac conducted in HCT116 90 by the ENCODE consortium, and were used for visualization. Peak and annotated genes can be found in Supplementary File 1.

### CAGE sequencing

To investigate the transcriptional consequences upon TBX3 overexpression, we performed Cap Analysis of Gene Expression followed by next generation sequencing (CAGE-seq). This method allows genome-wide transcriptional profiling of genes as well as accurate identification of transcriptional start sites (TSSs) and their corresponding promoter regions.

#### Library preparation and sequencing

HCT116 were transfected with either TBX3-3xFlag (donated by Peter J. Hurling) or the empty vector control, each in three biological replicates. Total RNA was isolated by using a standard TRIzol-based method (Cat#15596018, Thermo Scientific). The quality of the extracted RNA was tested on the Agilent 2100 Bioanalyzer. Each sample consisted of 5lzμg of initial total RNA. Library preparation, including cap-trapping step and polyadenylation, was conducted according to a standard CAGE protocol 31. Briefly, RNA was reverse transcribed using random primers, followed by biotinylation of the RNA cap and 3’ ends. Single-stranded, non-hybridized RNAs were digested with RNase leaving a 5’ cDNA that was captured by streptavidin beads. Sequencing was performed following the manufacturer instructions.

### Data Analysis

Quality of the obtained sequencing reads was assessed using FastQC (Andrews S. (2010; https://www.bioinformatics.babraham.ac.uk/projects/fastqc/). Bad quality (phred score < 33) and N containing reads were filtered out by fastx_trimmer (FASTX Toolkit, version 0.0.14) and removeN (Moirai), respectively. Reads matching tibosomal DNA (U13369.1.fa) were removed with RNAdust (version 1.06) and adapter sequence contaminants were removed with Trimmomatic 91, version 0.38, with the following settings: SE -phred33 LEADING:30 TRAILING:25 SLIDINGWINDOW:5:30 MINLEN:30. Trimmed and filtered reads were aligned to the UCSC human reference genome hg38 using the Burrows-Wheeler Aligner (BWA) 92, version 0.7.10-r789, with BWA-backtrack algorithm and the following settings: -n 0.02 -o 1 -e -1 -I 5 -d 10 -l 32 -k 2 -m 2000000 -M 3 -O 11 -E 4 - R 30 -q 0. Unmapped reads were re-aligned with Hisat2 93, version 2.1.0. Obtained alignment files in SAM format were converted to the binary BAM format, coordinate sorted and indexed with SAMtools 94, version 1.11-5-g0920974. Next, CAGE alignments were processed by PromoterPipeline ^95^ in order to get CTSS (CAGE transcription start sites) and CAGE peaks, which were associated with nearby Gencode v38 gene models by using ChIPseeker v1.32.0 package for R. CAGE peaks located within known promoter regions (2kbp) were used as training set in TSSClassifier ^96^ to predict whether sequence composition of remaining peaks is similar to promoters. CAGE peaks located in known promoters or classified as promoter associated were used for gene expression analysis and masked for enhancers calling ^97^. CAGE promoters to enhancers associations defined for pairs located within 500kbp and correlation test p.value < 0.05. CAGE expression profiles, promoters, enhancers, and associations are available through Zenbu browser: https://fantom.gsc.riken.jp/zenbu/gLyphs/#config=TBX3%20CAGE. CAGE regions, annotated genes, counts and differential expression can be found in Supplementary File 2.

### Integration of CAGE-seq and CUT&RUN

Genomic coordinates of enriched CAGE regions were input into a bed file and overlapped with filtered H3K27ac mediated chromatin loops, keeping loops with at least one anchor in a DEG region. Next, these loops were overlapped with TBX3 and then β-catenin ICEBERG peak sets. The corresponding CAGE/H3K27ac/TBX3/β-catenin loops were annotated to genes using GREAT, using coordinates of both anchors of each loop. Pathway enrichment was performed as described above. The results can be found in Supplementary File 3.

### BioID proximity labeling

#### BirA Plasmid construction

The BioID fused-protein construct was prepared by cloning the TBX3 gene sequence into a BioID pCS2 flag plasmid in-frame with the biotin ligase BirA at the C-terminal. In brief, the TBX3 coding sequence was PCR-amplified using a gradient-based PCR program (98°C 1 min – 98°C 30 sec – 62°C 30 sec – 72°C 1 min – GOTO Step 2 7x – 98°C 30 sec – 72°C 2 min – GOTO step 6 30x – 72°C 5 min – hold 4°C). PCR product was purified using MicroElute® Cycle-Pure-Kit (Omega Bio-Tek) and ligated into the BioID plasmid using type II restriction enzymes BamHI and XhoI (New England BioLabs) and T4 DNA ligase. 3 µL of ligation product was used for transformation of competent DH5α cells (Thermo Scientific) and plated on ampicillin-selective agar plates. 4 colonies were selected for further culturing before cell lysis and plasmid DNA purification using MiniPrep Kit (Thermo Scientific).

To generate the NPF motif deleted TBX3-BirA construct, the NEB Q5 Site-directed Mutagenesis Kit (New England BioLabs, USA) was used, following the manufacturer’s instructions. All constructs were confirmed by partial sequencing (Eurofins Genomics).

#### Cell transfection, pull-down and protein sample preparation

HEK293T cells were seeded in T175 to reach a 40% confluency. After 6 h, cells were transfected with 45 µg DNA (42 µg of either EV-BirA, TBX3-WT-BirA or TBX3-ΔNPF-BirA and 3µg empty vector GFP) using calcium phosphate. Fresh medium was added 18 h after transfection. At 42 h after transfection, the 20 mL culture media was supplemented with 0.1µM biotin (LS-3500, Iris Biotech), and either 0.01 µM CHIR99021 (Cat. #SML1046, Sigma Aldrich) or 0.02 µM LGK (Cat. # S7143, Selleck Chemicals). After an additional 24 h the cells were detached with Trypsin EDTA 0.25% (Cat# 25200056, Thermo Scientific) and subjected to three washed with PBS (Cat #14190094, Thermo Scientific).

Nuclear extraction was performed according to Schreiber et al., 1989. Briefly, cells were washed in 1 mL TBS and centrifugated at 1500 rpm. Cells were then resuspended in 400 µL buffer A (10 mM HEPES pH 7.9, 10 mM KCl, 0.1 mM EDTA, 0.1 mM EGTA, 1 mM DTT, 1mM PMSF (Cat# 36978, Thermo Fisher Scientific) and incubated at 4 °C for 15 min. Then, 25 µl 10% IGEPAL (Cat# J61055-AE, Thermo Fisher Scientific) was added to lyse cells and vortexed briefly. After centrifugation at 10 000g for 30 sec, the cytosolic fraction was discarded. The pellet containing the nuclear fraction was resuspended in 1 mL ice-cold buffer C (20 mM HEPES pH 7.9, 0.4 M NaCl, 1 mM EDTA, 1 mM EGTA, 1 mM DTT, 1mM PMSF), and incubated for 15 min at 4°C while vigorously shaking.

After nuclear extraction, pre-washed streptavidin-coated sepharose beads (GE17-5113-01, Sigma Aldrich) were added to the cell lysates and incubated for 3 h at 4°C with end-over-end rotation. Protein-bound beads underwent four washes in 1 mL wash buffer (50 mM ammonium bicarbonate in MilliQ) and were subsequently digested in 200 µL wash buffer with 0.4 µg (Pierce Trypsin Protease, Cat# PI90057, Thermo Scientific) for 1 hour at 37°C on an end-over-end rotator. The resulting supernatant was set aside, and the beads were washed twice with additional 100 µL wash buffer. All post-digestion supernatants were combined (total volume: 400uL).

To reduce the sample, 16 µL 100 mM DTT (Cat# FERR0862, Thermo Scientific) was added and incubated for 30 min at 37°C with end-over-end rotation, followed by alkylation through the addition of 46 µL 100 mM Iodoacetamide (Cat# NC0312634, Sigma Aldrich) for 45 min at room temperature in the dark. For complete digestion, the samples were treated with an additional 0.5 µg trypsin overnight at 37°C on an end-over-end rotator. Subsequently, samples were desalted using Pierce C18 Spin Columns (Cat# 89870, Thermo Fisher Scientific) in accordance with the manufacturer’s instructions, and then dried via speed vacuum centrifugation at room temperature.

#### Mass spectrometry data acquisition and analysis

10 µL of sample was transferred to Polypropylene Snap Top Microvials (6ERV11-08PPC, Thermo Fisher Scientific) and analyzed by mass spectrometry, using an Easy nano LC 1200 system interfaced with a nanoEasy spray ion source (Thermo Fisher Scientific) connected Q Exactive HF Hybrid Quadrupole-Orbitrap Mass Spectrometer (Thermo Fisher Scientific). The peptides were loaded on a pre-column (Acclaim PepMap 100, 75 μm x 2 cm, Thermo FisherScientific) and the chromatographic separation was performed using an EASY-Spray C18 reversed-phase nano LC column (PepMap RSLC C18, 2 μm, 100A 75 μm x 25 cm, Thermo Fisher Scientific). A linear gradient of 6-30% buffer B (0.1% formic acid in acetonitrile) against buffer A (0.1% formic acid in water) during 65 min and 100% buffer B against buffer A till 90 min was carried out with a constant flow rate of 300 nL/min. Separated peptides were electrosprayed and analyzed using a Q-Exactive HF mass spectrometer (Thermo Fisher Scientific), operated in positive polarity in a datadependent mode. Full scans were performed at 120,000 resolutions at a range of 380–1400 m/z. The top 15 most intense multiple charged ions were isolated (1.2 m/z isolation window) and fragmented at a resolution of 30,000 with a dynamic exclusion of 30.0 s.

Raw mass spectrometry data was analyzed for peptide identification and quantification with Proteome Discoverer 2.5 (Thermo Fisher Scientific), using the SequestHT search engine against the Homo sapiens UniProt database (UP000005640; 852,685 entries). Cysteine carbamidomethylation was used as static modification and methionine oxidation as dynamic modification for both identification and quantification. A maximum of 2 trypsin cleavages were allowed, and the precursor and fragment mass tolerance were 15 ppm and 0.1 Da, respectively. Peptides with false discovery rate (FDR) of less than 0.01 were considered significant, and the minimum peptide length was set to 5. Streptavidine and BirA were included in the database for exclusion in the search.

Processed mass spectrometry data were further analyzed to determine enrichment of proteins in treatment samples over controls using SAINTexpress ^98^, an implementation of the significance analysis of interactome (SAINT) algorithm ^99^. Total spectral counts were used as input. A SAINT score > 0.6 and an estimated Bayesian false discovery rate (BFDR) < 0.05 were used as thresholds for significance. Overlap analysis and plotting of data were performed with the R programming language (R Core Team, 2017, in Rstudio /Rstudio Team, 2015). The results of all tested conditions can be found in Supplementary File 4. Biological pathway enrichment analysis was performed with the online tool Enrichr ^100^, which can be found in Supplementary File 5.

### Western blot

Cultured HEK293T cells transfected with either TBX3-BirA or BirA-only expressing plasmid, in WNT ON and WNT OFF condition, were harvested and washed per BioID assay protocol. Samples were subjected to whole cell lysis, as well as subcellular fractionation lysis. For the latter, cells were resuspended in lysis buffer (PBS, 1% NP40, protease inhibitor) for two washes, followed by sonication (40 amplitude, 5 sec on, 5 sec off, 10 sec runtime), yielding cytoplasmic and nuclear fractions. 30 µL of each sample was treated with 24 µL Laemmli buffer and 6 µL Mercaptoethanol, and boiled at 95°C for 15 min. Samples were then subjected to SDS-PAGE separation on a pre-cast gel (BioRad) and blotting on a polyvinylidene difluoride (PVDF) membrane probed with rabbit monoclonal anti-histone H3 (Cat# 07690, EMD Millipore, 1:25000-50000), mouse monoclonal anti-alpha tubulin (Cat# NB100-690, Novus Biologicals, 1:5000) and mouse monoclonal anti-FLAG® M2 (Cat# F1804-50UG, Sigma Aldrich, 1:1000).

### Protein structure and protein complex prediction

Structural predictions were done with ColabFold (v.1.5.3), a software package that offers an integrated protein prediction solution as a webbased interface (utilizing Google Colaboratory notebooks). More specifically, the AlphaFold2.ipynb notebook was used ^101^. This tool integrates rapid homology searches, employing many-against-many sequence searching (MMseqS2) against the databases UniRef100, PDB100, and environmental sequence database, in conjunction with AlphaFold2 and RoseTTAFold methodologies ^102–104^.

The predictions were executed with the standard settings recommended by the developers. In brief, the input sequences (Supplementary File 11) were provided to the query_sequence command. Num_relax was set to 0. Template mode was set to none. When predicting complexes consisting of multiple proteins or domains the sequences of all subunits were concatenated using a colon (‘:’). MMseqS2_uniref_env, in unpaired_paired mode, was selected for multiple sequence alignment. For modeling, alphafold2_multimer_v3 was chosen for protein complexes, while alphafold2_ptm was selected for protein structure modeling. The default setting of 5 models with one single seed value was used. Recommended settings for both models were used to predict protein structure. Protein structure/ interactions were checked for consistency across five generated models. The results presented in the figures pertain solely to the top-ranking model (rank 1 out of 5). For monomer predictions the top-ranking model was determined by the highest average local distance difference test (LDDT) score—an indicator of model accuracy estimation. For prediction protein complexes the five generated models were ranked by their pTM and ipTM scores with the following formula: 0.2xpTm + 0.8xipTM. For monomer predictions standard T4 TPU’s with 12gb RAM were used. To model protein complexes, a faster A100 GPU with 16gb RAM was utilized.

The decision to exclusively generate models for the highly structured T-box domain was motivated by the presence of the NPF motif, while also the remainder of the protein exhibited predominantly unstructured characteristics. Similarly, more accurate predictions of protein complexes were attained by focusing solely on modeling conserved protein domains. In the context of modeling protein complexes and NPF interactions, instances were classified as ‘not interacting’ when domains were evidently distant from the NPF motif, clear prediction errors were discernible (such as intercalating structures), or distinct models exhibited significant divergence. The settings to generate Figure 6. C,D and Supplementary Figure 2. C,D,E were adapted from ^105^. Visualization of pdb files was done with Mol* ^106^ of RCSB PDB. All PDB files of the structural models can be found in Supplementary Files 6-10.

### Protein conservation prediction

Protein conservation estimates were done with CoservFold (V.1, 2023)(ConservFold.ipynb Colaboratory (google.com)) (Graham C, Stansfeld P and Rodrigues C, Conservation-Colab: Conservation to 3D structure, Github, 10.5281/zenodo.10062701, 2023). This Google Collaboratory based platform generates a multiple sequence alignment file with MMSEQ2 ^103^. Subsequent run of Weblogo3 ^107^ calculates entropy scores from the sequence similarities. The predictions were executed with the standard settings recommended by the developers. The TBX3 T-box protein sequence derived from previously published crystal structure available in the protein data bank (PDB) as accession number 1H6F was used as query sequence.

Weblogo: Information content in bits, overall height of character indicates positional conservation.

3D Structure: Conservation is indicated as the B factor that is calculated by Entropy2 for each amino acid. It is visualized with red being relatively high and blue being relatively low (95% confidence limits). It was visualized with PyMOL Molecular Graphics System, Version 2.6 Schrödinger.

### TOPFlash luciferase reporter assay

To conduct reporter assays, HEK293T cells were seeded in a 96-well plate overnight and co-transfected with 50 ng of firefly reporter, 5 ng of renilla control, and 10 ng of the plasmid of interest in each well. M50 Super 8x TOPFlash (Addgene Plasmid #12456) and M51 Super 8x FOPFlash (mutant, Addgene #12457) as used before ^26,108^. If necessary, cells were treated with control or CHIR99021 (1 µM, Cat. #SML1046, Sigma Aldrich) 6 h after transfection. The dual luciferase assay was performed with some modifications as previously described ^109^. After overnight incubation, cells were lysed in passive lysis buffer (25 mM Tris, 2 mM DTT, 2 mM EDTA, 10% (v/v) glycerol, 1% (v/v) Triton X-100, pH 7.8) and agitated for 10 min. The lysates were then transferred to a flatbottomed 96-well luminescence assay plate. Firefly luciferase buffer (200 μM D-luciferin in 200 mM Tris-HCl, 15 mM MgSO4, 100 μM EDTA, 1 mM ATP, 25 mM DTT, pH 8.0) was added to each well and the plate was incubated for 2 min at room temperature. The luciferase activity was measured using a SpectraMax iD3 Multi-Mode Microplate Reader (Molecular Devices). Subsequently, Renilla luciferase buffer (4 μM coelenterazine-h in 500 mM NaCl, 500 mM Na2SO4, 10 mM NaOAc, 15 mM EDTA, 25 mM sodium pyrophosphate, 50 μM phenyl-benzothiazole, pH 5.0) was added to the plate, and luminescence was immediately measured. The data were normalized to the Renilla control values, performed in triplicate, and the Top/Renilla ratio was used as an indicator of β-catenin-driven transcription.

### Statistical analysis

GraphPad Prism software was used for statistical analyses. Values were expressed as mean ± standard deviation. Mann Whitney U test was used to analyze the differences between two groups and p<0.05 was considered statistically significant. All experiments were performed at least three times and all samples analyzed in triplicate unless otherwise stated.

## Notes

### Competing Interest Statement

The authors have declared no competing interest.

