## Supplementary Figures for "The Developmental Transcription Factor TBX3 Physically Engages with the Wnt/β-catenin Transcriptional Complex in Human Colorectal Cancer Cells to Regulate Metastasis Genes"

Supplementary Figure 1 (for Figure 5)

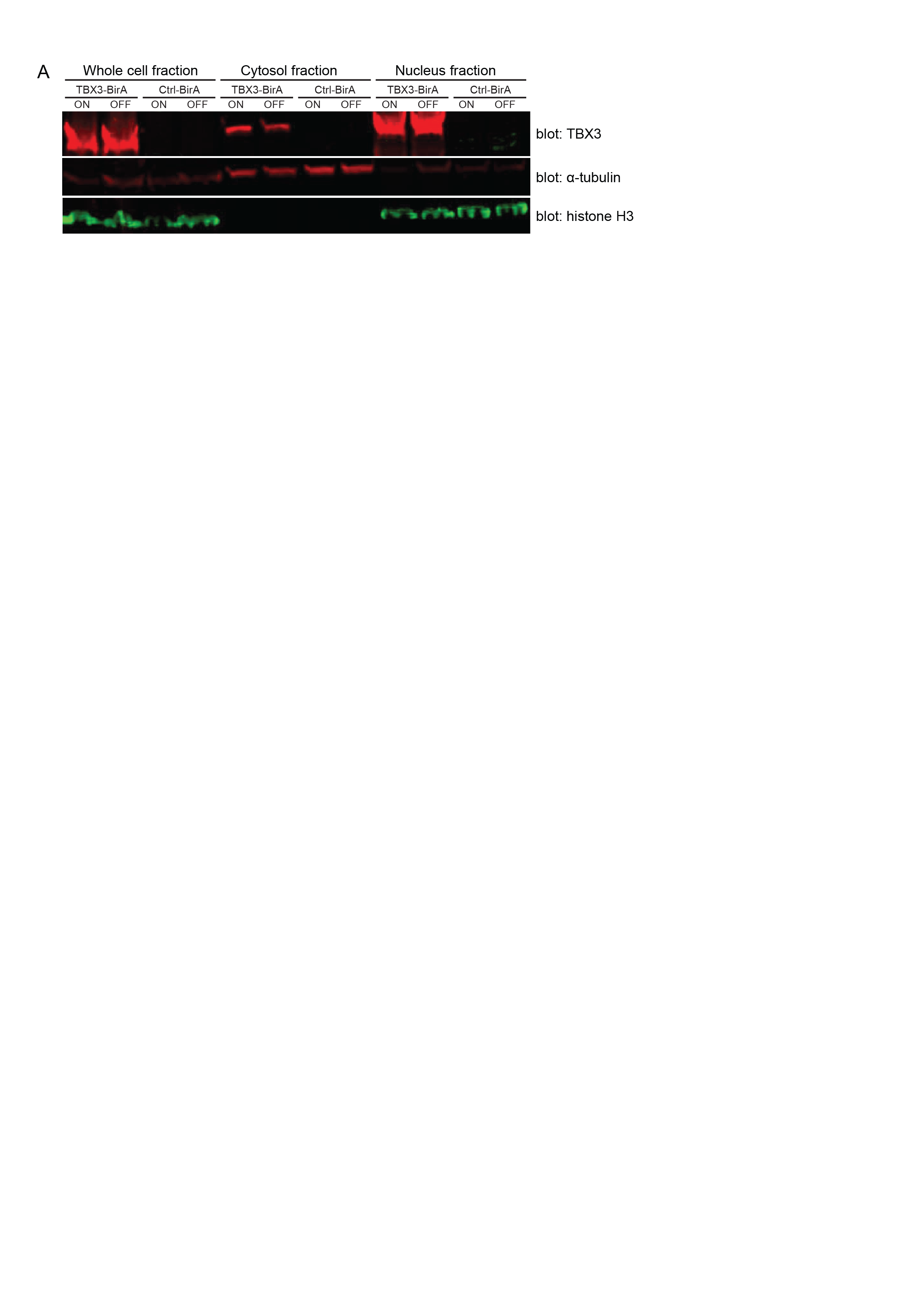

Supplementary Figure 1. Validation of BirA-fused TBX3 expression and nuclear localization.

(A) HEK293T were transfected with TBX-BirA and empty vector BirA (ctrl-BirA) and cultured in ‘WNT-ON’ and ‘WNT-OFF’ condition (ON, OFF, respectively). The whole cell, cytosolic and nuclear fractions were blotted against TBX3, α-tubulin and histone H3.

Supplementary Figure 2 (for Figure 6)

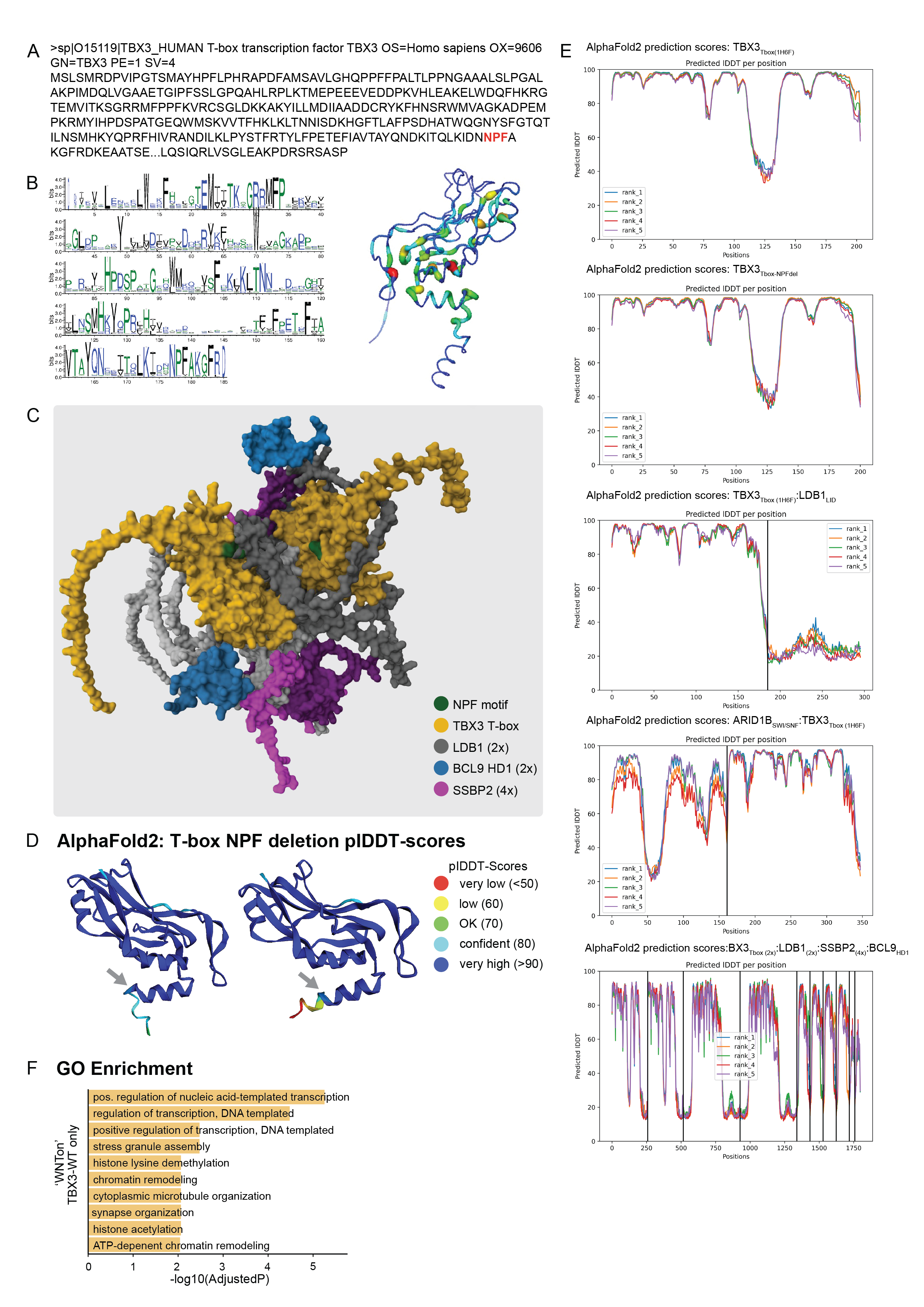

Supplementary Figure 2. (A) Amino acid sequence of human TBX3 with the NPF-motif highlighted in red.

(B) Left: WebLogo of the ConservFold prediction of the full T-box domain. Right: 3D model of T-box domain with the conservation score (=B-factor) colored on a scale from blue to red (blue = low, red = high).

(C) Multi-protein complex structure, to which we refer to as “Cthulhu”, starring as a AlphaFold prediction of the 3D structural composition of a protein complex containing 2x T-box domains (yellow, in green their NPF motifs), 2x full length LDB1 (gray), 4x full length SSBP2 (purple), 2x BCL9 HD1 domains (blue). The corresponding pdb file is provided as a Supplementary File (“Supplementary File 10.pdb”).

(D) AlphaFold2 3D structure prediction of TBX3 T-box (PDB 1H6F, left) and upon NPF deletion, (right). Color scheme indicates predicted IDDT values (pIDDT, model confidence out of 100).

(E) Confidence scores of the AlphaFold2 structural prediction of the sequence indicated above each plot. Y-axis: pIDDT. X-axis: protein sequence. Vertical lines separate different components of a multimer complex.

(F) Enrichment of GO terms among hits that are unique to the TBX3-WT vicinity in ‘WNT-ON’ condition only.
