## Supplementary material for "The Developmental Transcription Factor TBX3 Physically Engages with the Wnt/β-catenin Transcriptional Complex in Human Colorectal Cancer Cells to Regulate Metastasis Genes": Readme_Supplementary Files.docx

**Supplementary File 1:**

CUT&RUN data

**Supplementary File 2:**

TBX3 overexpression in HCT116 CAGE results: regions, annotations, counts, differentially expressed genes

**Supplementary File 3:**

KEGG enrichment results of list of genes resulting from CAGE data integration with CUT&RUN and HiChIP.

**Supplementary File 4:**

Table of BioID results for all tested conditions.

**Supplementary File 5:**

Table of BioID Enrichments results (GO, KEGG pathways and CORUM) for WNTon TBX3-WT only

**Supplementary File 6:**

PDB file of the TBX3 Tbox model.

TBX3Tbox1H6F_unrelaxed_rank1_alphafold2_ptm_model5_seed0

**Supplementary File 7:**

PDB file of the TBX3 Tbox model with an NPF deletion.

TBX3tboxdNPF_unrelaxed_rank1_alphafold2_ptm_model4_seed0

**Supplementary File 8:**

PDB file of the TBX3 Tbox and the LDB1 LID domain model.

TBX3tboxLDB1lid_unrelaxed_rank1_alphafold2_multimer_v3_model3_seed0

**Supplementary File 9:**

PDB file of the TBX3 Tbox and the ARID1B swi/snf model.

TBX3TBoxARID1Bswisnf_unrelaxed_rank1_alphafold2_multimer_v3_model2_seed0

**Supplementary File 10:**

PDB file of the whole complex (“Chtulu”) containing 2x TBX3 T-box, 2x LDB1, 4x SSDB2, 2x BCL9 HD1.

Chtulu_TBOX2xLDB12xSSBP4xBCL9HD12x_unrelaxed_rank1_alphafold2_multimer_v3_model4_seed0

**Supplementary File 11:**

List of Protein sequences used for structure predictions.
