## Supplementary material for "The Developmental Transcription Factor TBX3 Physically Engages with the Wnt/β-catenin Transcriptional Complex in Human Colorectal Cancer Cells to Regulate Metastasis Genes": Supplementary File 11.docx

**Protein sequences for AlphaFold2 (multimer) predictions**

**Sequences for Figure 6.B (PDB 1H6F dimer)**

>1H6F_1|**Chains A, B**|T-BOX TRANSCRIPTION FACTOR TBX3|HOMO SAPIENS (9606)

MKDDPKVHLEAKELWDQFHKRGTEMVITKSGRRMFPPFKVRCSGLDKKAKYILLMDIIAADDCRYKFHNSRWMVAGKADPEMPKRMYIHPDSPATGEQWMSKVVTFHKLKLTNNISDKHGFTILNSMHKYQPRFHIVRANDILKLPYSTFRTYLFPETEFIAVTAYQNDKITQLKIDNNPFAKGFRDTGNGRR

>1H6F_2|**Chains C, D**|5'-D(*TP*AP*AP*TP*TP*TP*CP*AP*CP*AP*CP*CP*TP* AP*GP*GP*TP*GP*TP*GP*AP*AP*AP*T)-3'|SYNTHETIC (32630)

TAATTTCACACCTAGGTGTGAAAT

**Sequences for Figure 6.C**

### Left: TBX3 (UniProt O15119) T-Box domain wild-type

DDPKVHLEAKELWDQFHKRGTEMVITKSGRRMFPPFKVRCSGLDKKAKYILLMDIIAADDCRYKFHNSRWMVAGKADPEMPKRMYIHPDSPATGEQWMSKVVTFHKLKLTNNISDKHGFTILNSMHKYQPRFHIVRANDILKLPYSTFRTYLFPETEFIAVTAYQNDKITQLKIDNNPFAKGFRD

**Right: TBX3 (UniProt O15119)** **T-Box domain - NPF deleted**

DDPKVHLEAKELWDQFHKRGTEMVITKSGRRMFPPFKVRCSGLDKKAKYILLMDIIAADDCRYKFHNSRWMVAGKADPEMPKRMYIHPDSPATGEQWMSKVVTFHKLKLTNNISDKHGFTILNSMHKYQPRFHIVRANDILKLPYSTFRTYLFPETEFIAVTAYQNDKITQLKIDNAKGFRD

**Sequences for Figure 6.D**

**Left: TBX3 (UniProt O15119) T-Box domain wild-type: ARID1B (UniProt Q8NPD5) SWI/SNF domain**

DDPKVHLEAKELWDQFHKRGTEMVITKSGRRMFPPFKVRCSGLDKKAKYILLMDIIAADDCRYKFHNSRWMVAGKADPEMPKRMYIHPDSPATGEQWMSKVVTFHKLKLTNNISDKHGFTILNSMHKYQPRFHIVRANDILKLPYSTFRTYLFPETEFIAVTAYQNDKITQLKIDNNPFAKGFRD:

SLAKRCICVSNIVRSLSFVPGNDAEMSKHPGLVLILGKLILLHHEHPERKRAPQTYEKEEDEDKGVACSKDEWWWDCLEVLRDNTLVTLANISGQLDLSAYTESICLPILDGLLHWMVCPSAEAQDPFPTVGPNSVLSPQRLVLETLCKLSIQDNNVDLIL

**Right: TBX3 (UniProt O15119) T-Box domain wild-type: _linker_ LDB1 (UniProt Q86U70) LID domain _linker_**

DDPKVHLEAKELWDQFHKRGTEMVITKSGRRMFPPFKVRCSGLDKKAKYILLMDIIAADDCRYKFHNSRWMVAGKADPEMPKRMYIHPDSPATGEQWMSKVVTFHKLKLTNNISDKHGFTILNSMHKYQPRFHIVRANDILKLPYSTFRTYLFPETEFIAVTAYQNDKITQLKIDNNPFAKGFRD:

_MSGGSTMSSGGGNTNNSNSKKKSPASTFALSSQVP_DVMVVGEPTLMGGEFGDEDERLITRLENTQFDAANGIDD_EDSFNNSPALGANSPWNSKPPSSQESKSENPTSQASQ_

**Supplementary Figure X.C**

**TBX3 (UniProt O15119) T-box (2x):LDB1(UniProt Q86U70, 2x): SSBP2 (UniProt 81877, 4x):BCL9(UniProt O00512) HD1**

DDPKVHLEAKELWDQFHKRGTEMVITKSGRRMFPPFKVRCSGLDKKAKYILLMDIIAADDCRYKFHNSRWMVAGKADPEMPKRMYIHPDSPATGEQWMSKVVTFHKLKLTNNISDKHGFTILNSMHKYQPRFHIVRANDILKLPYSTFRTYLFPETEFIAVTAYQNDKITQLKIDNNPFAKGFRD:

DDPKVHLEAKELWDQFHKRGTEMVITKSGRRMFPPFKVRCSGLDKKAKYILLMDIIAADDCRYKFHNSRWMVAGKADPEMPKRMYIHPDSPATGEQWMSKVVTFHKLKLTNNISDKHGFTILNSMHKYQPRFHIVRANDILKLPYSTFRTYLFPETEFIAVTAYQNDKITQLKIDNNPFAKGFRD:

MSVGCACPGCSSKSFKLYSPKEPPNGNAFPPFHPGTMLDRDVGPTPMYPPTYLEPGIGRHTPYGNQTDYRIFELNKRLQNWTEECDNLWWDAFTTEFFEDDAMLTITFCLEDGPKRYTIGRTLIPRYFRSIFEGGATELYYVLKHPKEAFHSNFVSLDCDQGSMVTQHGKPMFTQVCVEGRLYLEFMFDDMMRIKTWHFSIRQHRELIPRSILAMHAQDPQMLDQLSKNITRCGLSNSTLNYLRLCVILEPMQELMSRHKTYSLSPRDCLKTCLFQKWQRMVAPPAEPTRQQPSKRRKRKMSGGSTMSSGGGNTNNSNSKKKSPASTFALSSQVPDVMVVGEPTLMGGEFGDEDERLITRLENTQFDAANGIDDEDSFNNSPALGANSPWNSKPPSSQESKSENPTSQASQ:

MSVGCACPGCSSKSFKLYSPKEPPNGNAFPPFHPGTMLDRDVGPTPMYPPTYLEPGIGRHTPYGNQTDYRIFELNKRLQNWTEECDNLWWDAFTTEFFEDDAMLTITFCLEDGPKRYTIGRTLIPRYFRSIFEGGATELYYVLKHPKEAFHSNFVSLDCDQGSMVTQHGKPMFTQVCVEGRLYLEFMFDDMMRIKTWHFSIRQHRELIPRSILAMHAQDPQMLDQLSKNITRCGLSNSTLNYLRLCVILEPMQELMSRHKTYSLSPRDCLKTCLFQKWQRMVAPPAEPTRQQPSKRRKRKMSGGSTMSSGGGNTNNSNSKKKSPASTFALSSQVPDVMVVGEPTLMGGEFGDEDERLITRLENTQFDAANGIDDEDSFNNSPALGANSPWNSKPPSSQESKSENPTSQASQ:

GMYGKGKSNSSAVPSDSQAREKLALYVYEYLLHVGAQKSAQTFLSEIRWEKNITLGEPPGFLHSWWCVFWDLYCAAPERRETCEHSSEAKAFHDY:

GMYGKGKSNSSAVPSDSQAREKLALYVYEYLLHVGAQKSAQTFLSEIRWEKNITLGEPPGFLHSWWCVFWDLYCAAPERRETCEHSSEAKAFHDY: GMYGKGKSNSSAVPSDSQAREKLALYVYEYLLHVGAQKSAQTFLSEIRWEKNITLGEPPGFLHSWWCVFWDLYCAAPERRETCEHSSEAKAFHDY: GMYGKGKSNSSAVPSDSQAREKLALYVYEYLLHVGAQKSAQTFLSEIRWEKNITLGEPPGFLHSWWCVFWDLYCAAPERRETCEHSSEAKAFHDY:

AMAAKVVYVFSTEMANKAAEAVLKGQVETIVSFHI:

AMAAKVVYVFSTEMANKAAEAVLKGQVETIVSFHI
